# A chimeric Ad5-Envp-VLP vaccine platform confers broad-spectrum immunity against emerging and re-emerging pathogens

**DOI:** 10.1101/2025.07.03.663092

**Authors:** Yuan Zhang, Caiqian Wang, Yanhong Zheng, Feiyu Chen, Yaya Feng, Lingying Fang, Zongmei Wang, Zhen F. Fu, Ming Zhou, Ling Zhao

## Abstract

Integrating complementary vaccine modalities is essential for combating emerging infectious threats. Here, we developed Ad5-Envp-VLP, a chimeric adenoviral platform synergizing adenoviral delivery efficiency with virus-like particle (VLP) structural mimicry. This system stably produces self-assembling VLPs in suspension HEK293 cultures, exhibiting enhanced immunogenicity over soluble antigens. Following intramuscular immunization, the platform induces early B cell expansion and sustains germinal center reactions, driven by the upregulation of B cell cycle-related genes (*Cdc6*, *Cdc45*, *Cdc20*, *Cdc25C*, *Aurka*, *Aurkb*, and *Ccnb1/2*) and robust T follicular helper (Tfh) cell differentiation, generating durable neutralizing antibodies against both influenza virus and rabies virus. These effects are conserved across mouse, canine, and feline models. Crucially, integrated flow cytometry and scRNA-seq demonstrate that intranasal delivery recruits and functionally reprograms lung innate immune cells (notably alveolar macrophages and dendritic cells), driving mucosal sIgA secretion and CTL responses. A single nasal dose confers lasting protection against homologous and heterologous influenza A strains. The platform also elicits cross-neutralizing antibodies against SARS-CoV-2 variants. Together, Ad5-Envp-VLP thus establishes a modular vaccine platform for antigenically plastic pathogens by combining *in vivo* self-assembly with dual pulmonary-muscular delivery.

**One Sentence Summary:** Novel mucosal and systemic vaccine delivers self-assembling particles, triggering strong lung immunity and broad virus protection across species.

## INTRODUCTION

The global emergence of SARS-CoV-2 and persistent antigenic drift in influenza viruses have exposed critical limitations in conventional vaccine paradigms, particularly their inability to concurrently achieve single-dose durability, broad cross-protection, and thermostability(*1*). Virus-like particles (VLPs), represent a transformative advance, retaining native virion structure without replication risks while enabling robust B cell receptor (BCR) clustering and multi-epitope T cell activation(*2, 3*). Commercial VLP vaccines, such as Gardasil® 9 for HPV and Recombivax HB® for hepatitis B, have reduced the incidence of their target diseases by more than 90% globally (*4, 5*). Nevertheless, conventional VLP platforms face three key challenges: (1) manufacturing complex arising from precise 3D assembly requirements and costly purification workflows(*6*); (2) dependence on adjuvant co-formulation and multi-dose regimens to overcome poor immunogenicity(*7*); and (3) thermolability necessitating stringent cold-chain logistics (e.g., Gardasil®9’s 2–8°C storage), which severely restricts its deployment in resource-limited regions. These challenges underscore the urgent need for next-generation VLP systems that integrate scalable production, intrinsic adjuvant properties, and thermal resilience while maintaining the integrity of the antigen structure.

Recent advances in mRNA vaccine engineering enabled *in vivo* self-assembly of VLPs, combining the antigen design flexibility and rapid adaptability of mRNA platforms with the multivalent antigen presentation of VLPs(*8*). The RQ3013-VLP candidate exemplified this paradigm, utilizing mRNAs encoding SARS-CoV-2 spike (S), membrane (M), and envelope (E) proteins to drive spontaneous VLP formation *in vivo*. This approach elicits superior neutralizing antibody (NAb) titers and T-cell responses compared to conventional spike-only mRNA formulations(*9*). Further optimization strategies include engineering chimeric S proteins with heterologous cytoplasmic tails (e.g., HIV-1/SIV Gag) to enhance VLP assembly efficiency, broadening neutralizing antibody breadth against SARS-CoV-2 variants(*10*). The S-EABR (ESCRT- and ALIX-binding region) system represents another breakthrough, exploiting host ESCRT (Endosomal Sorting Complex Required for Transport) machinery to drive spike protein self-assembly into enveloped VLPs (eVLP)(*11*). This “encoding-expression-assembly” integrated technology bypasses the cumbersome *in vitro* VLP production, forming dense and ordered antigen arrays *in vivo* with only a single component without introducing heterogeneous antigens. Despite these innovations in VLP assembly *in vivo* based on the mRNA platform, clinical translation remains constrained because intramuscular administration prioritizes systemic immunity over mucosal protection in respiratory tissues(*12–14*). While aerosolized mRNA-LNP formulations show promise for pulmonary delivery, their requirements for multiple dose and thermolability limit distribution in resource-limited regions(*15*).

Recombinant adenoviral vectors (rAd) have emerged as an ideal gene delivery platform due to intrinsic safety features (replication-deficient design), high transfection efficiency, and scalable production(*16*). Notably, their capacity for aerosolized inhalation delivery enables direct activation of respiratory mucosal immunity, offering unique advantages for vaccine development against influenza viruses, coronaviruses, and other respiratory pathogens(*17*). Preclinical validation of inhaled Ad5-nCoV showed that it induces durable lung-resident T cell memory, activates both mucosal and systemic immunity, and possesses systemic reactogenicity (febrile incidence: <5% vs. 28% for intramuscular administration), illustrating a “single-dose, dual-protection” paradigm(*18*). Structural analyses reveal that Spike proteins expressed by Ad5-nCoV maintain native-like glycosylation patterns and conformational epitopes, enabling potent neutralization antibody responses against emerging variants(*19*). In addition, our previous study based on adenoviral vector engineering has shown that dual insertion of foreign genes into the E1/E3 region enhanced target gene expression and immunogenicity(*20*). VLPs primarily activate humoral immunity through repetitive antigen display but lack endogenous adjuvant properties inherent to adenoviral vectors. Although adenoviral vectors and VLPs have complementary advantages in immune activation and multivalent display of antigens, their integration for enhanced vaccine immunogenicity remains underexplored.

In this study, we developed Ad5-Envp-VLP, a novel recombinant adenoviral platform that combines the advantages of VLP and adenoviral vectors using the EABR strategy—a strategy capable of spontaneously forming Envp-VLPs in animals. Using the hemagglutinin (HA) protein of influenza A virus (IAV) as a model and immunizing via two immunization routes, intranasal and intramuscular injection, we evaluated the effectiveness of Ad5-Envp-VLP in mice. We found that VLPs formation significantly enhanced vaccine immunogenicity through distinct immune mechanisms depending on the administration route. For the intramuscular route, the Ad5-Envp-VLP rapidly activated B cells, promoted germinal center formation and maintenance, and thereby elicited potent and durable humoral immune responses. Following intranasal administration, Ad5-Envp-VLP recruited and activated innate immune cells (e.g., dendritic cells, macrophages) in lungs, significantly enhancing antigen processing and presentation capabilities across multiple cell types. Ad5-Envp-VLP also elevated the migratory capacity and metabolic activity of innate immune cells. This process ultimately promoted the generation of a higher level of antigen-specific IgA antibodies and facilitated the production of tissue-resident memory T cells (TRM) in the lung. Importantly, Ad5-HA_PR8_-VLP-induced local pulmonary mucosal immunity provided mice with long-lasting protection against multiple heterologous influenza viruses. Additionally, we validated the broad applicability of Ad5-Envp-VLP against diverse pathogens and across species using SARS-CoV-2 and rabies virus models. In summary, this platform combines the high transduction efficiency of adenoviral vectors with the multivalent antigen display of VLPs, creating a versatile framework for developing next-generation vaccines against evolving zoonotic pathogens.

## RESULTS

### Self-assembled HA-EABR chimeric VLPs via rAd vector induce potent antibody responses

Previous study employing an mRNA-based platform demonstrated that incorporating the EABR motif into the cytoplasmic tail of the SARS-CoV-2 spike protein facilitates recruitment of the host ESCRT machinery, enabling self-assembly of spike-enveloped VLPs (S-eVLPs) through membrane budding processes(*11*). To evaluate whether this strategy could be generalized to recombinant adenoviral platforms, we fused the EABR motif to influenza A hemagglutinin proteins (HA, residues 1–552). The chimeric construct (HA-EABR) was cloned into recombinant adenoviral genomes to generate Ad5-HA_PR8_-VLP. Parental adenovirus (Ad5-HA_PR8_) encoding full-length envelope protein served as controls (Fig. 1A). Both constructs displayed comparable replication kinetics in HEK293 cells (Fig. 1B). Western blot analysis confirmed HA secretion in supernatants of Ad5-HA_PR8_-VLP-infected cells, with intracellular HA levels reduced by approximately 50% compared to Ad5-HA_PR8_-infected cells (Fig. 1C), indicating enhanced EABR-mediated budding. To assess cross-species compatibility, MDCK (canine), BSR (murine), and CRFK (feline) cells were infected (MOI=1) with Ad5-eGFP, Ad5-HA_PR8_, or Ad5-HA_PR8_-VLP. Consistently, HA protein was detected in supernatants of Ad5-HA_PR8_-VLP-infected cells across all species, accompanied by respective intracellular HA reductions of 50% (MDCK), 30% (BSR), and 35% (CRFK) (Fig. S1). To confirm whether the HA protein detected in the supernatant existed in the form of VLPs, we separated the protein from the virus in the supernatant by iodixanol gradient density centrifugation, and the separated protein layer was further purified by size exclusion chromatography (SEC). Transmission electron microscopy (TEM) of SEC-purified material revealed spherical vesicles (90–110 nm in diameter) densely decorated with surface protrusions (Fig. 1D).

**Figure 1.**
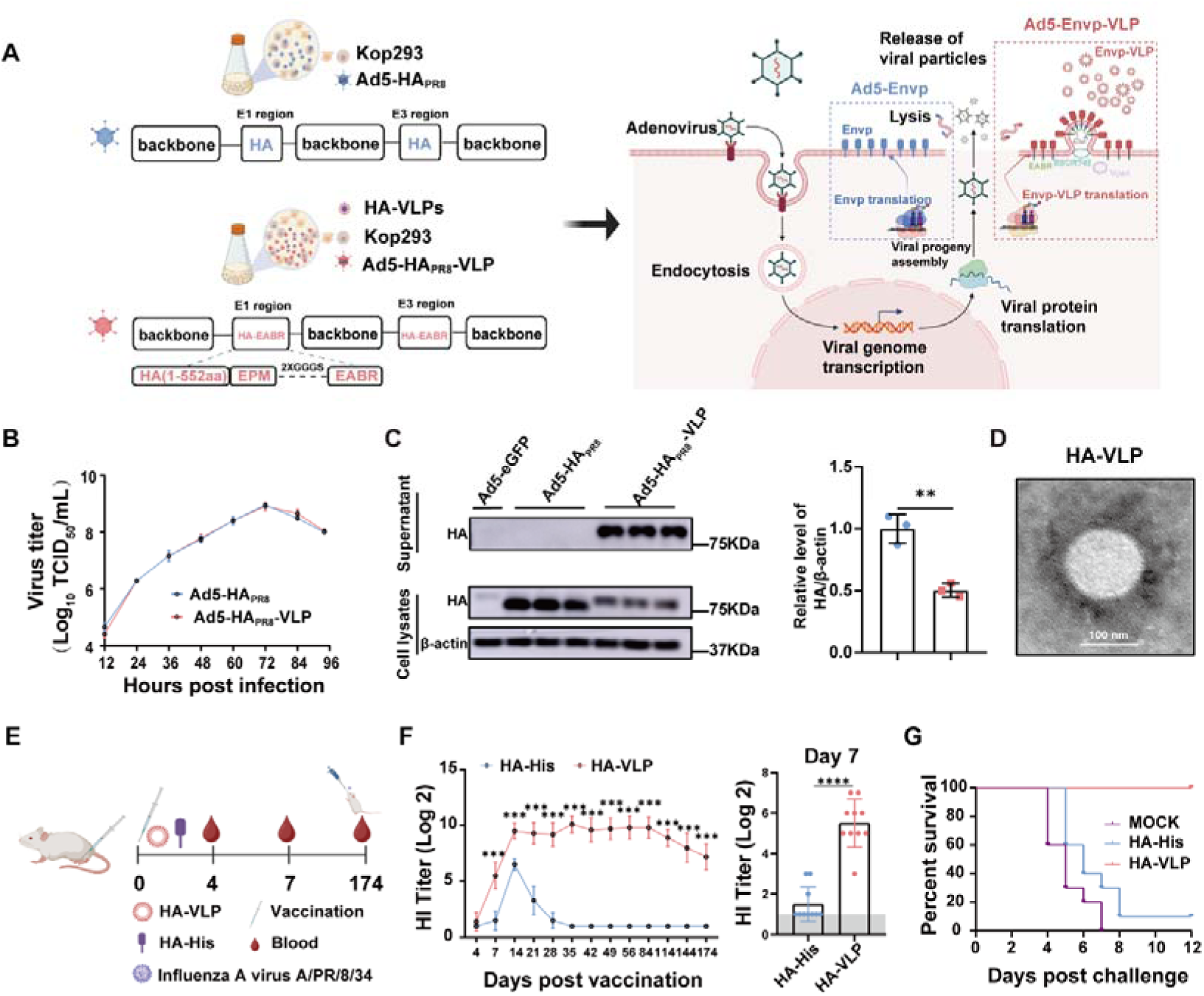
Construction and characterization of Ad5-HA_PR8_ and Ad5-HA_PR8_-VLP. (A) Schematic design of Ad5-HA_PR8_ and Ad5-HA_PR8_-VLP. The EABR motif was fused to cytoplasmic domain-truncated HA (residues 1–552). Parental adenovirus Ad5-HA_PR8_ encoding full-length HA protein served as controls. (B) Replication kinetics of Ad5-HA_PR8_ and Ad5-HA_PR8_-VLP in HEK 293 cells (n=3). (C) Western blot analysis of HA protein expression in cell lysates and supernatants from Ad5-HA_PR8_-VLP-infected cells versus controls. (D) Transmission electron microscopy of SEC-purified VLPs from Ad5-HA_PR8_-VLP-infected cells, revealing spherical particles (90 nm) with surface projections. Scale bars: 100 nm. (E) Immunization and challenge protocol. Female BALB/c mice (6-week-old, n=10/group) were immunized intramuscularly with DMEM, 10 μg HA-VLP, or HA-His (AS03-adjuvanted), followed by intranasal challenge with 10□ PFU PR8 (H1N1) at 174 dpi. Serum HI titers were measured at indicated intervals. (F) HI titers during monitoring (left) and at 7 dpi (right). (G) Survival rates monitored for 12 days post-infection. Data are mean ± SD. Statistical analysis: unpaired two-tailed t-test (C), two-way ANOVA with Tukey’s test (F), and log-rank test (G). NS: not significant; \**P* < 0.05, \*\**P* < 0.01, \*\*\**P* < 0.005, \*\*\*\**P* < 0.0001.

Compared with soluble subunit vaccines, VLP-based vaccines often exhibit enhanced immunogenicity due to their dense antigen display and structural mimicry of native virions(*21*). To explore the immunogenicity of purified VLPs, mice were immunized with 10 μg (quantified by ELISA) purified HA-VLP or soluble HA protein (HA-His) adjuvanted with AS03 (Fig. 1E). Immunogenicity of both antigens was evaluated by HI assay. At 7 dpi, seroconversion rates in the HA-VLP group reached 100%, while HI activity was detected in only 30% of the HA-His-immunized mice (Fig. 1F, right). While HA-His immunized mice reached peak HI titers (GMT=2^6^) at day 14 followed by rapid decline to undetectable levels by day 35, the HA-VLP group showed higher and more persistent HI titer throughout the monitoring period (Fig. 1F, left). Consistent with the antibody kinetics, HA-VLP immunization conferred complete protection (100% survival) against lethal intranasal challenge with 10^4^ PFU of PR8, whereas HA-His immunization resulted in 90% mortality (Fig. 1G). In conclusion, these results demonstrated that EABR-modified HA protein expressed by adenovirus vector self-assembled into VLPs in infected multi-species cells, and the formed VLPs elicited rapid and durable neutralizing antibody responses conferring long-term protection against viral challenge.

### Ad5-HA_PR8_-VLP elicits robust humoral immunity via early B cell activation and sustained germinal centers responses

To assess the impact of VLPs formation on recombinant adenoviral vaccine efficacy, we compared the immunogenicity of Ad5-HA_PR8_ and Ad5-HA_PR8_-VLP in BALB/c mice. Mice were intramuscularly immunized in the hind limb with 10□ TCID_50_/100 μL of either construct (Fig. 2A). At 4 days post-immunization (dpi), HI antibodies were detected in 40% of Ad5-HA_PR8_-VLP-immunized mice, whereas no measurable antibody response was observed in the Ad5-HA_PR8_ group (Fig. 2B, left). ELISA analysis revealed a 3.95-fold increase in HA-specific IgM titers in the Ad5-HA_PR8_-VLP group (1:522) compared to the Ad5-HA_PR8_ group (1:132) (Fig. 2D), suggesting that the VLPs produced *in vivo* accelerates early humoral immune responses initiation. At 7 and 14 dpi, Ad5-HA_PR8_-VLP elicited significantly higher HI titers (Fig. 2B, right) and HA-specific IgG titers than Ad5-HA_PR8_ (Fig. 2C). HI antibody titers gradually declined in both groups during the 174-day immune monitoring period (Fig. 2B, left). Intravital imaging revealed exogenous expression at the injection site for approximately 25 days following intramuscular administration of 10^7^ TCID_50_ Ad5-Luci (Fig. S2A), suggesting that sustained antigen availability may underlie the long-lasting humoral immunity induced by adenoviral vectors.

**Figure 2.**
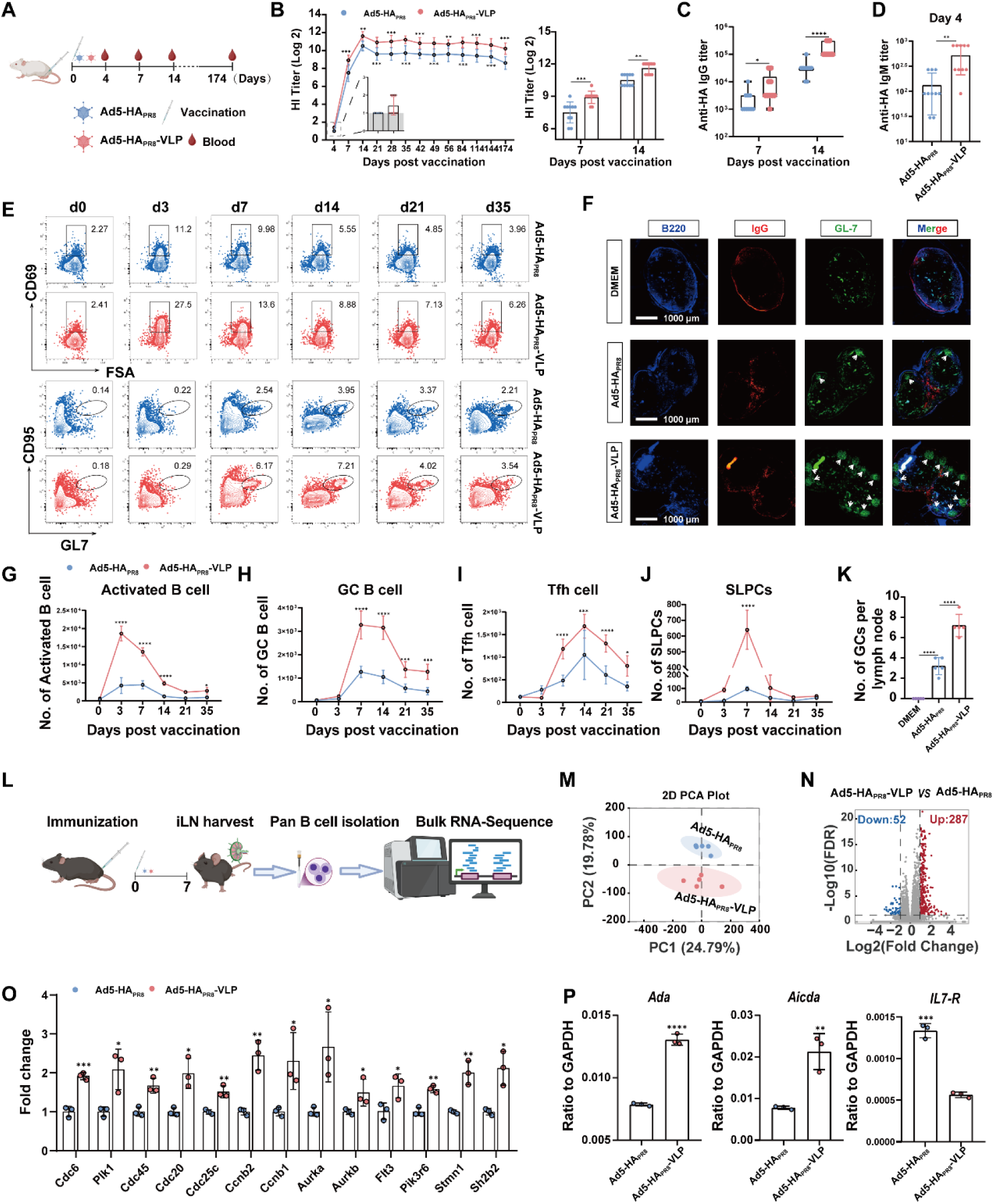
Enhanced humoral immunity and B cell activation driven by Ad5-HA_PR8_-VLP immunization. (A) Experimental design: BALB/c mice (n=10/group) were intramuscularly immunized with 10^7^ TCID_50_/100 μL of Ad5-HA_PR8_ or Ad5-HA_PR8_-VLP. Serum samples were harvested at indicated time-point. (B) Hemagglutination-inhibiting (HI) antibody kinetics. Left: Longitudinal HI titers over 174 days and seroconversion rates at day 4; Right: HI titers at days 7 and 14 post-immunization. (C) HA-specific IgG titers at days 7 and 14. (D) HA-specific IgM titers at day 4. (E) Representative FCM plots of activated B cells (CD45+B220+CD3-CD69+, top) and GC B cells (CD45+B220+CD3-CD95+GL7+) in iLNs. (F) Representative immunofluorescence results of GC formation in inguinal lymph nodes at day 7 (B220+, blue; IgG+ red; GL7+, green;). Scale bars, 1000 μm (leftmost column only). (G-J) The statistical results of activated B cell (G), GC B cell (H), Tfh cell (CD45+B220-CD3+PD-1+CXCR5+, I) and short-lived plasma cells (SLPCs, CD45+B220+CD3-CD138+CD44+, J) in iLNs. (K) Quantification of GC numbers per iLN section. (L) Schematic of experimental design. C57BL/6 mice (n=5 per group) were intramuscularly immunized with Ad5-HA_PR8_-VLP or Ad5-HA_PR8_. iLNs were collected at 7-day post-immunization for pan-B cell isolation and bulk RNA sequencing. (M) Principal component analysis (PCA) plot demonstrating distinct clustering between VLP-formulation and control groups. (N) Volcano plot of differentially expressed genes (DEGs). Red: upregulated genes (FDR <0.01, log2[fold change] >1); blue: downregulated genes (FDR <0.01, log2[fold change] <-1); gray: nonsignificant genes. (O) qPCR validation of *PIK3-AKT* and *Cell cycles pathway* related genes. (P) The *Ada*, *Aicda* and *IL7R* mRNA level were quantified by qPCR. Data presentation: Mean ± SD. Statistical analysis: (B, C, G-J and O) two-way ANOVA with Tukey’s test; (D, K and P) unpaired two-tailed t-test. \**P* < 0.05, \*\**P* < 0.01, \*\*\**P* < 0.005, \*\*\*\**P* < 0.0001.

To explore the mechanism by which Ad5-HA_PR8_-VLP induces early and high-titer antibody production, we systematically explored the B cell activation kinetics after immunization with the two constructs (Ad5-HA_PR8_ and Ad5-HA_PR8_-VLP). FCM results revealed that the number of activated B cells (CD69+) increased by 4.39-fold on day 3 after Ad5-HA_PR8_-VLP immunization than that of Ad5-HA_PR8_ (Fig. 2E top and G), indicating that Ad5-HA_PR8_-VLP promoted early B cell activation. These activated B cells exhibited preferential migration to lymphoid follicles for germinal center (GC) initiation, supported by follicular helper T (Tfh) cells(*22*). Dynamic tracking of GC B and Tfh cells demonstrated that Ad5-HA_PR8_-VLP induced a stronger and more sustained GC reaction from day 7 to 35 post-immunization than that of Ad5-HA_PR8_ (Fig. 2E bottom, H and I), indicating that Ad5-HA_PR8_-VLP induced a robust and sustained germinal center response. In addition, immunofluorescence staining of iLNs at day 7 post-immunization showed a 2.25-fold increase in GC numbers in the Ad5-HA_PR8_-VLP group compared to Ad5-HA_PR8_ (Fig. 2F and K), suggesting that the produced VLPs promoted a robust germinal center response in the early stage of immunization. B cells failing to enter GCs were directly activated in T cell zones via T cell-independent (TI) or partially T cell-dependent (TD) pathways, differentiating into short-lived plasma cells (SLPCs). We found the Ad5-HA_PR8_-VLP group exhibited a 6.6-fold higher SLPC count at the peak response (day 7) than the Ad5-HA_PR8_ group (Fig. 2J), suggesting that Ad5-HA_PR8_-VLP induced early SLPC differentiation to produce antibodies more rapidly. Notably, the Ad5-HA_PR8_-VLP immunized group generated significantly more HA-specific GC B cells, MBCs (Fig. S3A and B), and long-lived plasma cells (LLPCs) than Ad5-HA_PR8_ group (Fig. S3C). Collectively, our results suggest that Ad5-HA_PR8_-VLPs induces high-titer neutralizing antibody production via early activation of B cells and potent germinal center responses.

### Ad5-HA_PR8_-VLPs induce B cell transcriptional reprogramming in lymph nodes

To elucidate the mechanisms underlying VLPs-induced enhancement of B cell activation, maturation, and differentiation in lymph nodes, we performed transcriptomic profiling of B cells isolated from mice immunized with Ad5-HA_PR8_ or Ad5-HA_PR8_-VLP at 7 dpi (Fig. 2L). PCA revealed distinct clustering of transcriptomic profiles between groups, indicating significant differences in the gene expression patterns of B cells driven by VLPs and membrane antigens (Fig. 2M). Differential expression analysis identified 339 DEGs (287 upregulated, 52 downregulated) in pan-B cells from Ad5-HA_PR8_-VLP compared with the Ad5-HA_PR8_ group reprogramming (Fig. 2N), indicating that VLPs perturbed transcriptional reprogramming of B cells in iLNs. KEGG pathway enrichment highlighted four dominant pathways: *Cell cycle*, *Cytokine-cytokine receptor interaction*, *Endocytosis*, and *PI3K-AKT signaling* (Fig. S4A). Consistent with KEGG, GO analysis demonstrated enrichment of mitosis-related biological processes (*chromosome segregation, nuclear division, mitotic nuclear division, spindle organization*) (Fig. S4B), which further indicate that VLPs driven B cell proliferation through mitotic activation. Critically, we observed pronounced upregulation of *Aicda* (Activation-Induced Cytidine Deaminase), a master regulator of somatic hypermutation (SHM) and class-switch recombination (CSR). This result was further validated by qPCR, where Ad5-HA_PR8_-VLP promoted *Aicda* transcription (Fig. 2P), implying that sustained VLPs exposure promoted antibody diversification by facilitating genomic remodeling in B cells. Moreover, we found that Ad5-HA_PR8_-VLP promoted B cell activation pathways, inducing a significant upregulation of *Sh2b2* (2.1-fold, BCR signaling amplifier), *Ada* (1.9-fold, adenosine metabolism regulator), and *Stmn1* (2.0-fold, microtubule dynamics controller) compared to the Ad5-HA_PR8_ group. Transcript levels of *IL7R* (a marker of naive B cells) were significantly reduced following Ad5-HA_PR8_-VLP immunization (Fig. 2O and 2P), consistent with an increased proportion of mature B cells within the total B cell population. The coordinated upregulation of mitotic regulators (*Cdc6*, *Cdc45*, *Cdc20*, *and Cdc25c*) and cyclin-dependent kinases (*Aurka*, *Aurkb*, and *Ccnb1/2*) (Fig. 2O), combined with *Aicda*-mediated genomic instability pathways, implied VLPs’ dual functionality: driving clonal expansion through enhanced cell cycle progression, while enabling antibody maturation via SHM and CSR. Taken together, our results suggest that Ad5-HA_PR8_-VLP drives B cell transcriptional reprogramming in lymph nodes via secreted VLPs.

### Ad5-HA_PR8_-VLP enhances T cell activation and cytokine-polarized responses

Naïve T cell activation is initiated by antigenic peptide recognition via T cell receptors (TCRs) upon interaction with antigen-presenting cells (APCs). Prior studies identified B cells as the primary APCs driving naïve CD4+ T cell activation following nanoparticle-antigen immunization(*23*). Considering the effects of Ad5-HA_PR8_-VLP on B cell activation, we further evaluated the T cell response triggered by Ad5-HA_PR8_-VLP. The results showed that the CD69+ cell population in CD4+ and CD8+ T cells of Ad5-HA_PR8_-VLP-immunized mice was significantly increased at 7 dpi, compared with the Ad5-HA_PR8_ control group (Fig. S5A and B). Moreover, the Ad5-HA_PR8_-VLP group exhibited higher frequencies of effector memory T cells (TEM, CD62L−CD44+) in CD4+ (Fig. S5C, top) and CD8+ subsets (Fig. S5E, top), along with increased CD4+ central memory T cells (TCM, CD62L+CD44+) (Fig. S5C, bottom), indicating enhanced memory T-cell differentiation and subset polarization. In sum, these findings indicate that VLPs robustly activate T cells, enhancing both effector and memory T cell responses. To assess functional polarization, we quantified cytokine-producing HA-specific T cells via intracellular staining. At day 7, the Ad5-HA_PR8_-VLP group showed higher frequencies of IFN-γ+ single-positive, TNF-α+ single-positive, TNF-α+IL-2+IFN-γ− double-positive, and IL-2+ single-positive CD4+ T cells (Fig. S5D, top). By day 21, the Ad5-HA_PR8_-VLP group predominantly generated TNF-α+ single-positive and TNF-α+IL-2+IFN-γ− CD4+ T cells, whereas TNF-α+IL-2+ double-positive cells predominated in the Ad5-HA_PR8_ group (Fig. S5D, bottom). Notably, Ad5-HA_PR8_-VLP immunization induced 1.32-fold and 3.32-fold more TNF-α+ CD4+ T cells than Ad5-HA_PR8_ at days 7 and 21, respectively.

As key cytotoxic effectors of adaptive immunity, cytotoxic T lymphocytes (CTLs) exhibited divergent clonal expansion dynamics across distinct vaccine formulations. At day 7, while IFN-γ+ single-positive CD8+ T cells constituted the predominant subset in both Ad5-HA_PR8_-VLP and Ad5-HA_PR8_ groups, the Ad5-HA_PR8_-VLP group exhibited a 1.8-fold higher cell count (Fig. S5F, top). By day 21, Ad5-HA_PR8_-VLP induced substantial expansion of IFN-γ+TNF-α+IL-2− (3.72-fold), IFN-γ+TNF-α+IL-2+ (3.78-fold), and IFN-γ-TNF-α+IL-2+ (2.15-fold) CD8+ subsets versus Ad5-HA_PR8_ group (Fig. S5F, bottom). Strikingly, TNF-α+ single-positive CD8+ T cells were exclusively amplified in the VLP group (4.8-fold increase). Multifunctional CD8+ T cells (producing ≥2 cytokines) were 3.28-fold more abundant in VLP-immunized mice, underscoring enhanced polyfunctionality. Collectively, Ad5-HA_PR8_-VLP immunization drives robust T cell activation, promotes memory differentiation (TEM/TCM), and elicits TNF-α-biased CD4+ responses alongside polyfunctional CTL expansion.

### Ad5-HA_PR8_-VLP induced robust mucosal immune by enhancing lung T and B cell activation and differentiation

Adenoviral vectors are promising mucosal vaccine candidates because they effectively generate targeted immune protection at mucosal surfaces(*24*). To evaluate whether VLPs enhance adenoviral vaccine-induced mucosal immunity, BALB/c mice were intranasally immunized with 10□ TCID□□ of Ad5-HA_PR8_ or Ad5-HA_PR8_-VLP, and antibody titers were subsequently measured in serum and BALF at indicated time points (Fig. 3A). The serum antibody results showed that HI antibodies were detectable in both immunization groups at 14 dpi, but the titers in the Ad5-HA_PR8_-VLP group were significantly higher, which peaked at 28 dpi and then gently declined (Fig. 3B). Despite parallel decline kinetics, HI titers in Ad5-HA_PR8_-VLP-immunized group remained consistently higher than controls, underscoring sustained humoral efficacy. While peak HI antibody responses were uniformly achieved at day 28, one Ad5-HA_PR8_-immunized mouse retained baseline titers throughout the 174-day study (Fig. 3B, right), suggesting antigen-specific hyporesponsiveness. In addition to the systemic humoral immune response, the Ad5-HA_PR8_-VLP group demonstrated significantly elevated mucosal IgA titers in the BALF at 14 and 28 dpi, indicating that Ad5-HA_PR8_-VLP can induce potent mucosal immunity (Fig. 3C). *In vivo* imaging showed transient luciferase expression (approximately 10 days) in lungs following intranasal Ad5-Luci administration (Fig. S2B), a shorter duration than intramuscular immunization, presumably due to the rapid turnover of lung epithelial cells. Moreover, ELISA quantitative analysis showed that the HA concentration in the BALF of the Ad5-HA_PR8_-VLP group was 36 times that of the Ad5-HA_PR8_ group on day 4 (Fig. 3D). These results suggest that EABR motif-driven VLP secretion can significantly facilitate antigen availability in the lung mucosa.

**Figure 3.**
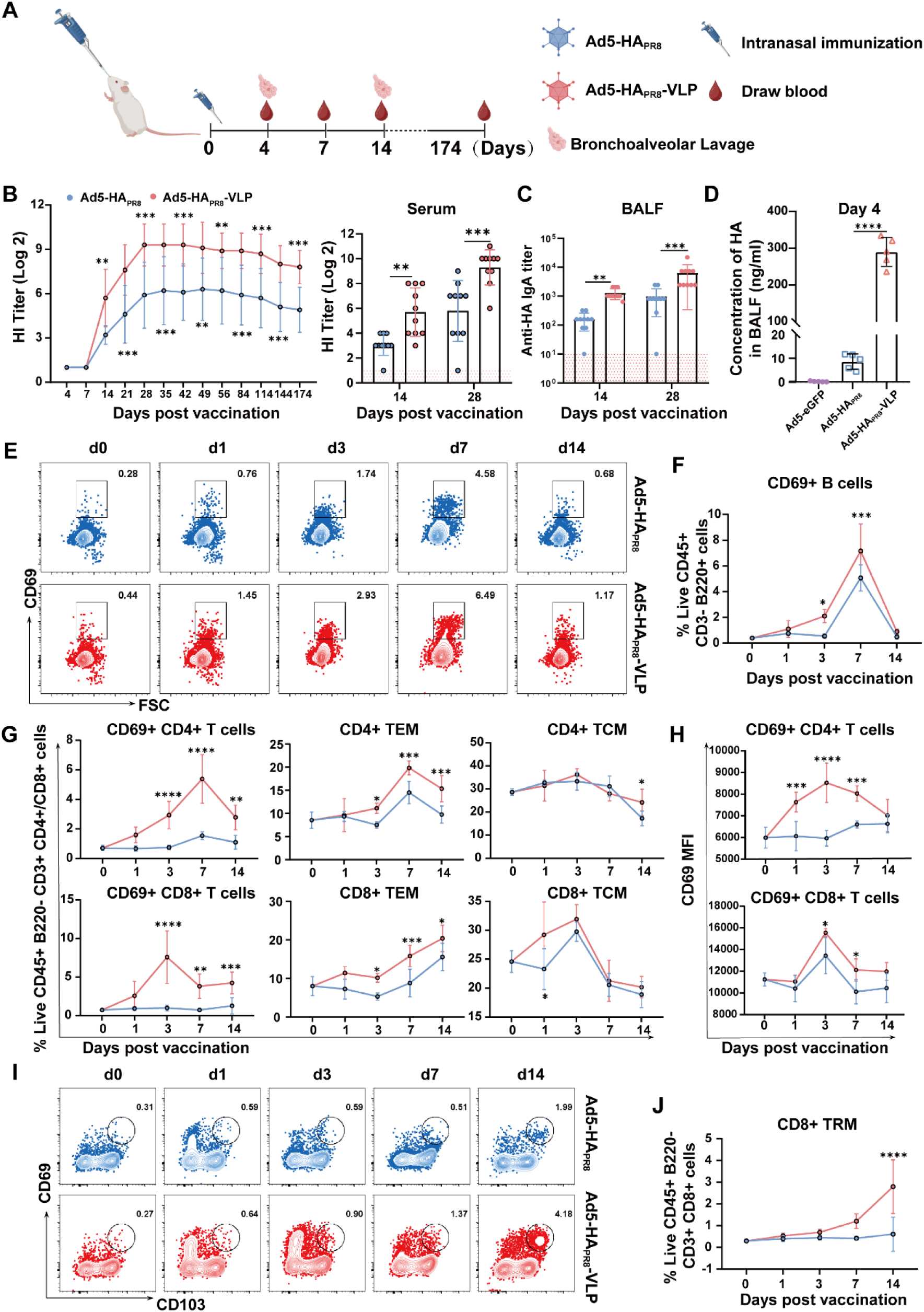
Ad5-HA_PR8_-VLP induces robust pulmonary mucosal immunity. (A) Experimental design schematic. BALB/c mice were intranasally immunized with 10□ TCID□□ Ad5-HA_PR8_-VLP or Ad5-HA_PR8_. Serum and BALF were collected at designated time points. (B) HI titers in serum across groups (n=10 per group) at indicated time points (left); HI titers at days 14 and 28 (right). (C) HA-specific IgA titers in BALF (n=10 per group) at days 14 and 28. (D) HA protein concentration in BALF (n=5 per group) at day 4. (E) Representative FCM plots of CD69+ B cells (CD45+CD3-B220+CD69+) in lungs at different time points (F) Quantification of CD69+ B cells in lungs across groups (n=5 per group). (G) Proportions of CD69+ T cells, effector memory T cells (TEM: CD44+CD62L-), and central memory T cells (TCM: CD44+CD62L+) in lungs (gated on CD45+B220-CD3+CD8-CD4+ or CD45+B220-CD3+CD8+CD4-). (H) MFI of CD69 in pulmonary T cells. (I) Representative FCM plots of lung tissue-resident memory CD8+ T cells (TRM, CD45+B220-CD3+CD8+CD69+CD103+) at indicated time points. (J) Proportions of CD8+ TRM in lungs (n=5 per group). Data are presented as mean ± SD. Statistical significance (Panels B, C, F, H, J) was determined by two-way ANOVA with Tukey’s multiple-comparison test; Panel D was analyzed by one-way ANOVA with Tukey’s test. Significance levels: NS (not significant), \**P* < 0.05, \*\**P* < 0.01, \*\*\**P* < 0.005, \*\*\*\**P* < 0.0001.

To assess B cell activation dynamics in the lung, we monitored CD69+ B cell proportions at multiple time points. As shown in Fig. 3E and F, CD69+ B cells in the VLP group expanded dynamically, starting from day 3, peaking on day 7, and sustaining higher levels than the control group through day 7. Meanwhile, the Ad5-HA_PR8_-VLP group exhibited higher proportions of both total IgA+ B cells and HA-specific IgA+ B cells compared to the Ad5-HA_PR8_ group at 21 dpi (Fig. S6A-C). Consistent with this finding, the data of ELISpot assay showed a significantly increased number of HA-specific IgA ASCs in the Ad5-HA_PR8_-VLP group (Fig. S6G and H). Interestingly, both total and HA-specific IgG2b+ B cells were observed exclusively in the Ad5-HA_PR8_-VLP group, with total IgG2b+ B cells showing a 2.8-fold higher proportion than the Ad5-HA_PR8_ group (Fig. S6D-F). In contrast, there were no significant differences in IgM+ B cells and IgG1+ B cells between the two groups at 21 dpi (Fig. S6I).

Parallel analysis of T cell activation revealed that CD4+ and CD8+ (CD69+) T cells in the Ad5-HA_PR8_-VLP group began to increase at 1 dpi, peaking at day 7 and day 3. While CD69+ CD4+ T cells in the Ad5-HA_PR8_ group also increased by day 7, their frequency was substantially lower than in the Ad5-HA_PR8_-VLP group (Fig. 3G). The frequency of CD69+ CD8+ T cells remained unchanged in the Ad5-HA_PR8_ group, though the normalized mean fluorescence intensity (MFI) of CD69 transiently upregulated at 3 dpi (Fig. 3G and H). Furthermore, differentiation analysis showed that both CD4+ and CD8+ TEM enriched in the Ad5-HA_PR8_-VLP group at days 3, 7, and 14, whereas TCM showed similar trends between groups (Fig. 3G). Intracellular cytokine staining further demonstrated that the Ad5-HA_PR8_-VLP group induced more TNF-α+IL-2+IFN-γ− CD4+ and CD8+ T cells than the Ad5-HA_PR8_ group (Fig. S6J). Only the Ad5-HA_PR8_-VLP group generated TNF-α+IL-2−IFN-γ+ CD4+ T cells and IFN-γ+ CD8+ T cells (Fig. S6J). Critically, CD8+ tissue-resident memory T cells (TRM, CD69+CD103+) expanded significantly in the Ad5-HA_PR8_-VLP group by day 14, whereas the Ad5-HA_PR8_ group showed no TRM increase (Fig. 3I and J), indicating that VLPs uniquely promote CD8+ TRM differentiation. In conclusion, Ad5-HA_PR8_-VLP significantly enhanced mucosal IgA titers, and IgG2-biased B cell responses, while eliciting rapid CD69+ B and T cell activation and differentiation.

### VLPs promotes pulmonary innate immune cell recruitment, activation, and maturation

Our data above showed delayed kinetics in HI antibody production (7-day lag) and B cell activation (4-day lag) following intranasal compared to intramuscular immunization. This temporal disparity suggests unique innate immune coordination in lung tissues during early antigen recognition. To elucidate this mechanism, we characterized innate immune cell dynamics in lung post-intranasal immunization (Fig. 4A). Lung DCs subsets exhibited differential responses. CD11b+ cDC proportions progressively increased from day 1 to 14 in Ad5-HA_PR8_-VLP-immunized mice, whereas Ad5-HA_PR8_ controls showed delayed expansion (initiated at day 7) with consistently lower proportions (Fig. 4B and C). Notably, while CD86 expression (a maturation marker) in CD11b+ cDC followed similar temporal trends between groups, Ad5-HA_PR8_-VLP immunization induced significantly higher CD86 levels than Ad5-HA_PR8_, indicating VLP-driven enhancement of CD11b+ cDC recruitment and maturation (Fig. 4D, top). In parallel, Ad5-HA_PR8_-VLP immunization triggered an immediate surge in CD103+ cDC proportions by day 1. Although both groups showed declined CD103+ cDC percentages by day 3, Ad5-HA_PR8_-VLP maintained a significantly higher proportion than Ad5-HA_PR8_ (Fig. 4C). Mirroring CD11b+ cDCs, CD86 expression in CD103+ cDCs peaked earlier and reached higher levels in the Ad5-HA_PR8_-VLP group (Fig. 4D, bottom), further supporting VLPs’ capacity to promote CD103+ cDC recruitment and functional maturation. Plasmacytoid dendritic cells (pDCs), a minor DC subset, play a critical role in antiviral immunity through robust type I interferon (IFN-I) secretion(*25*). Our data revealed no significant changes in pulmonary pDC proportions following either Ad5-HA_PR8_-VLP or Ad5-HA_PR8_ immunization (Fig. 4E). This finding suggests that VLPs production may not modulate pDC recruitment. As pivotal components of the pulmonary immune system, alveolar macrophages (AMs) play dual roles in host defense and immunomodulation(*26*). Both immunization groups showed initial AM depletion at day 1. However, Ad5-HA_PR8_-VLP immunized mice demonstrated rapid AM recovery by day 3, enabling AM proportions to remain significantly higher than those in Ad5-HA_PR8_ controls through day 14 (Fig. 4E). CD86 expression in AMs showed pronounced upregulation in the VLP group versus controls (Fig. 4F, left). Further analysis revealed distinct temporal patterns in lung F4/80+ macrophage responses between immunization groups. The proportion of F4/80+ macrophages exhibited a sustained elevation in the Ad5-HA_PR8_-VLP group, with a significant increase at day 3 (1.3-fold versus baseline) that persisted through day 14. In contrast, the Ad5-HA_PR8_ group showed only a transient elevation at day 1 (1.2-fold versus baseline), with fluctuating proportions thereafter (Fig. 4E). Furthermore, CD86 expression in the Ad5-HA_PR8_-VLP group showed progressive enhancement, with significant increases at both day 3 (1.4-fold versus baseline) and day 14 (1.5-fold versus baseline), while no CD86 upregulation was detected in Ad5-HA_PR8_ group (Fig. 4F, right). These coordinated patterns of cellular expansion and activation marker expression indicate that VLPs formulation drives robust, sustained macrophage activation compared to conventional adenoviral vectors.

**Figure 4.**
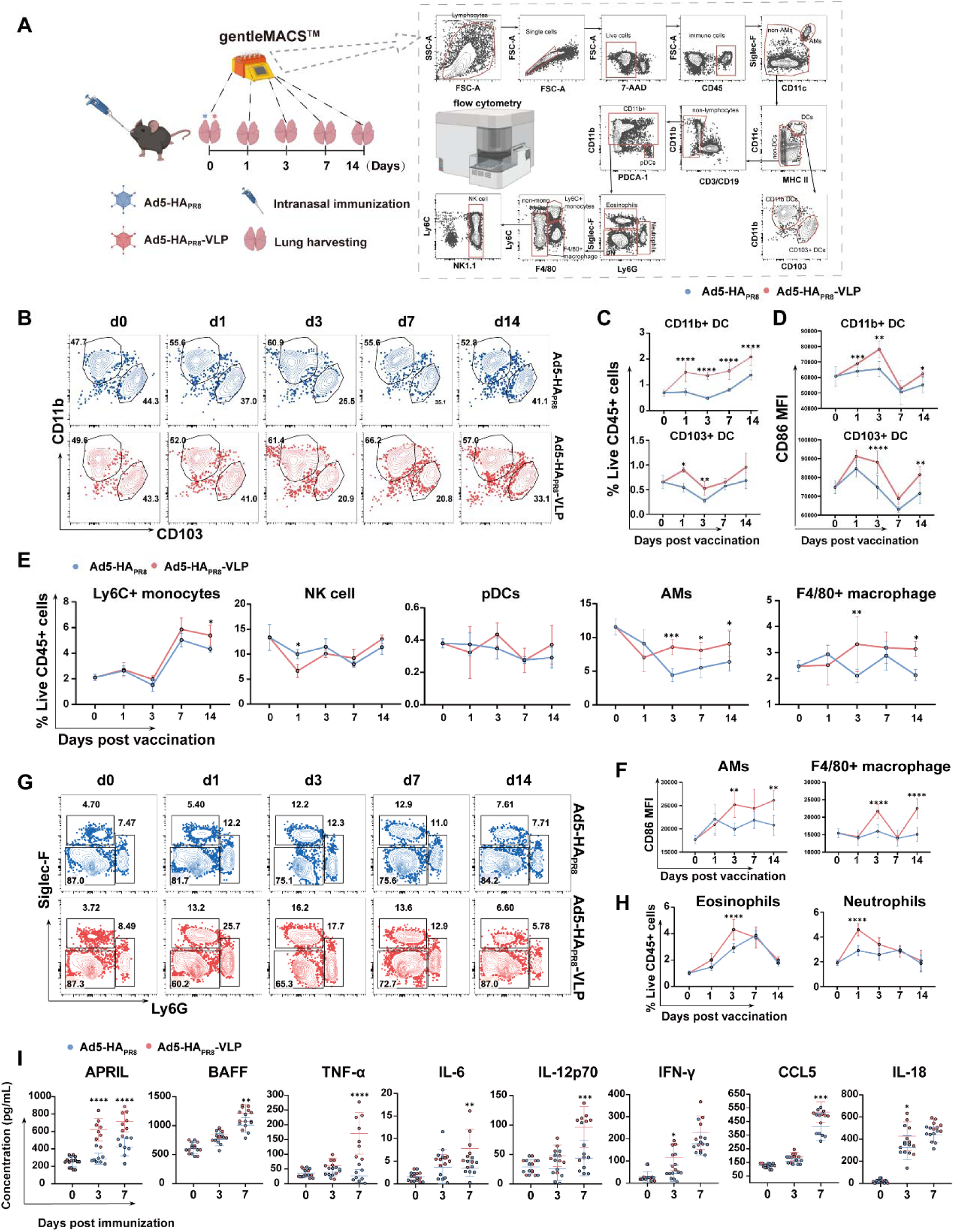
VLPs formulation promotes pulmonary innate immune cell recruitment, activation, and maturation. C57BL/6 mice (n=5 per group) were intranasally immunized with 10□ TCID□□ Ad5-HA_PR8_-VLP or Ad5-HA_PR8_. Lungs were harvested at indicated time points for analysis of innate immune cell populations. (A) Experimental design and FCM gating strategy for pulmonary innate immune cells. (B) Representative FCM plots of CD11b+ and CD103+ DCs in lungs. (C, D) Frequencies and CD86 MFI of CD11b+ and CD103+ DCs. (E) Frequencies of Ly6C+ monocytes, NK cells, pDCs, AMs, and F4/80+ macrophages. (F) CD86 MFI of AMs and F4/80+ macrophages. (G, H) Representative FCM plots and quantification of eosinophils and neutrophils in lungs. (I) Concentrations of cytokines in the lungs (n=8) at 3 and 7 dpi. Data are presented as mean ± SD. Statistical significance was determined by two-way ANOVA with Tukey’s multiple-comparison test. Significance levels: NS (not significant), \**P* < 0.05, \*\**P* < 0.01, \*\*\**P* < 0.005, \*\*\*\**P* < 0.0001.

NK cell dynamics revealed distinct early-phase suppression. Both groups showed significant NK cell depletion at day 1 (Ad5-HA_PR8_-VLP: 0.49-fold of baseline levels; Ad5-HA_PR8_: 0.75-fold of baseline levels), with more pronounced reduction in the Ad5-HA_PR8_-VLP group. Frequencies were normalized by day 14 without intergroup differences (Ad5-HA_PR8_-VLP: 0.97-fold; Ad5-HA_PR8_: 0.85-fold) (Fig. 4E). The proportion of Ly6C+ monocytes exhibited delayed expansion kinetics, initiated at day 7. Notably, the Ad5-HA_PR8_-VLP group showed significantly greater Ly6C+ monocyte proportions compared to Ad5-HA_PR8_ group by day 14 (Fig. 5E). Granulocyte responses showed distinct temporal patterns. The Ad5-HA_PR8_-VLP group exhibited a rapid eosinophil surge in lung tissue, peaking at day 3 with significantly higher counts than the Ad5-HA_PR8_ group, followed by a progressive decline through day 14. In contrast, the Ad5-HA_PR8_ group showed delayed accumulation that peaked at day 7 (Fig. 4G and H, left). Neutrophil infiltration exhibited formulation-dependent kinetics, with the Ad5-HA_PR8_-VLP group demonstrating a rapid 1.58-fold increase compared to Ad5-HA_PR8_ at day 1. This early surge was followed by progressive decline to baseline levels by day 14, indicating that VLP formulation preferentially enhances acute-phase neutrophil recruitment rather than sustaining long-term pulmonary infiltration (Fig. 4G and H, right). At day 3, concentrations of IL-18 and IFN-γ were significantly higher in the Ad5-HA_PR8_-VLP group than in the Ad5-HA_PR8_ group (Fig. 4I). This early elevation in IL-18 (a potent inflammasome product and IFN-γ inducer) and IFN-γ (a key activator of macrophages) suggests enhanced initial innate immune cell activation, particularly involving macrophages or NK cells. By day 7, the Ad5-HA_PR8_-VLP group demonstrated significantly elevated concentrations of APRIL, BAFF, TNF-α, IL-6, IL-12p70, and CCL5. This broader response indicates sustained and amplified inflammation (TNF-α, IL-6), enhanced Th1 priming (IL-12p70), robust B-cell activation signals (APRIL, BAFF), and increased chemotactic potential for immune cell recruitment (CCL5). In summary, these results demonstrate that VLPs enhance the recruitment, activation, and functional maturation of innate immune cell populations, thereby establishing a complex and well-orchestrated pulmonary immune microenvironment essential for initiating adaptive immunity.

**Figure 5.**
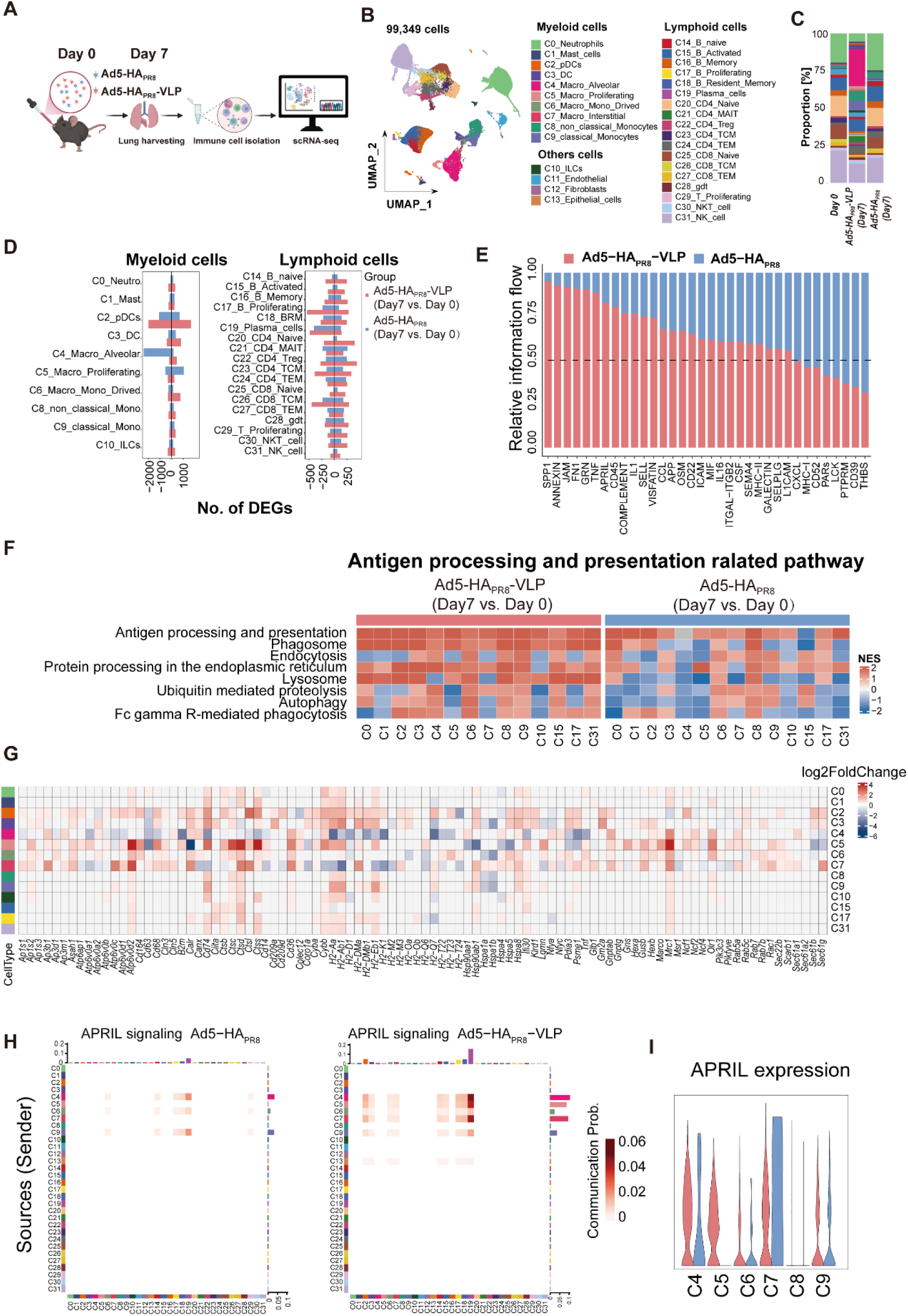
scRNA-seq revealed Ad5-HA_PR8_-VLP-driven remodeling of the pulmonary immune landscape. (A) Experimental design: C57BL/6 mice (n=3 per group) were intranasally immunized with 10□ TCID□□ Ad5-HA_PR8_-VLP or Ad5-HA_PR8_. Lungs were harvested at days 0 and 7 for scRNA-seq analysis of immune cell populations. (B) UMAP projection of 99,349 total lung cells clustered from scRNA-seq. (C) Proportion of immune cell subsets in different immune groups. (D) DEGs induced by Ad5-HA_PR8_-VLP and Ad5-HA_PR8_ immunization at day 7. (E) Relative information flow at 7 dpi. (F) Significantly enriched antigen processing and presentation KEGG pathways across clusters (C0-1, C15, C17 and C31) at day 7. Only clusters with a significantly modulated pathway are shown. (G) Heat map of key antigen processing and presentation response after Ad5-HA_PR8_-VLP immunization. (H) Heat map of APRIL signaling cell communication interactions in each immune group. (I) Violin plot of expression distribution of genes related to APRIL signaling pathway.

### ScRNA-seq revealed VLPs-enhanced pulmonary immune remodeling

To delineate VLPs-induced remodeling of the pulmonary immune landscape, we performed single-cell RNA sequencing (scRNA-seq) on lung immune cells at 0 and 7 dpi with Ad5-HA_PR8_-VLP and Ad5-HA_PR8_ (Fig. 5A). We captured 99,349 lung cells and annotated 31 distinct clusters encompassing immune cells (myeloid cells and lymphoid cells) and non-immune cells (lung endothelial cells, alveolar fibroblasts, epithelial cells) (Fig. 5B and Fig. S7A). Consistent with FCM results, Ad5-HA_PR8_-VLP vaccination significantly expanded APCs proportions, notably DCs, macrophages, and monocytes (Fig. 5C). Differential expression analysis demonstrated more DEGs in Ad5-HA_PR8_-VLP-vaccinated mice than in Ad5-HA_PR8_ controls, particularly within monocyte, macrophages, DC subsets, NK cells, T cells, and B cells (Fig. 5D). Gene set enrichment analysis (GSEA) showed robust enrichment of phagosome or lysosome formation and antigen presentation pathways across multiple cell types, especially pronounced in DCs (C2 and C3), AMs (C4), classical monocytes (C9), and activated B cells (C15) from Ad5-HA_PR8_-VLP-vaccinated mice (Fig. 5F). Additionally, we found that Ad5-HA_PR8_-VLP elicited significantly higher expression of key antigen processing and presentation genes (*CD74, Ctsb, Ctss, H2-Aa, H2-Ab1, H2-DMa, H2-Eb1, Hspa8, ifi30*) across immune cell clusters C2, C3, C5-C9, and C15 (Fig. 5G). Besides, Ad5-HA_PR8_-VLP specifically elevated *SEC61B* and *SEC61G* transcripts in DC clusters C2 and C3 (Fig. 5G), which have been shown to stabilize ER-endosome channels for antigen retro-translocation to the cytosol and subsequent MHC-I loading(*27*). SEC61G also regulates glycolysis and oxidative phosphorylation(*28*). GSEA of DCs (C2, C3) in the Ad5-HA_PR8_-VLP group showed significant positive enrichment of ‘Oxidative phosphorylation’ (NES=2.89, FDR=0), ‘Glycolysis’ (NES=1.53, FDR=0.03), and ‘Citrate cycle (TCA cycle)’ (NES=1.82, FDR=0) (Fig. S7B), indicating that VLPs enhance antigen presentation and energy metabolism in lung DCs. In macrophages, Ad5-HA_PR8_-VLP immunization significantly upregulated *NCF1* (p47-phox) and *NCF2* (p67-phox) – key cytoplasmic subunits of the NADPH oxidase complex regulating reactive oxygen species (ROS) generation(*29*) – in subsets C4 and C5 (Fig. 5G). CD36 mediates oxidized low-density lipoprotein (oxLDL) uptake, enhancing mitochondrial fatty acid oxidation (FAO) to amplify ROS burst and inflammation(*30*). Our data indicated that CD36 transcription increased in all macrophage clusters (C4-C7) (Fig.5G). The specific oxLDL receptor OLR1 was also upregulated in macrophages (C4-C6), suggesting VLPs enhance oxLDL uptake. OLR1 has been shown to amplify inflammatory cascades via NF-κB and MAPK pathways(*31*). Under hypoxic conditions, OLR1 and HIF-1α synergistically activate NLRP3 inflammasome-mediated inflammation. Accordingly, NF-κB, TNF, MAPK, and HIF-α signaling pathways were significantly enriched in macrophages after Ad5-HA_PR8_-VLP immunization (Fig. S7B).

Analysis of intercellular communication revealed increased information flow in pathways mediating immune cell migration and adhesion (SPP1, ANNEXIN, JAM, FN1, SELL, ICAM) after Ad5-HA_PR8_-VLP immunization (Fig. 5E and Fig. S7C). Pathways related to migration and adhesion were enriched in clusters C4-C6, C8, C9, C15, C17, C18, C22, C24, C27, C29, and C31 (Fig. S7B). Consistent with elevated lung APRIL protein levels (Fig. 4I), *APRIL* gene expression increased significantly in macrophages (C4-C7) (Fig. 5I), enhancing communication with proliferating B cells (C17), tissue-resident B cells (C18), and plasma cells (C19) (Fig. 5H), suggesting VLPs promote APRIL secretion in macrophage to enhance B cell proliferation and survival. Moreover, inflammation-related IL1 signaling flow was enhanced, with *IL18* specifically increased in DCs (C3) and macrophages (C5-C7) (Fig. S7D). While NK cell proportions decreased post-immunization in both groups, Ad5-HA_PR8_-VLP immunization induced significant positive enrichment of antigen processing and presentation, migration and adhesion pathways in NK cell (Fig. 5E and Fig. S7B). Pseudotime trajectory analysis of T and B cells showed Ad5-HA_PR8_-VLP immunization significantly decreased naïve T cell proportions (C20, C25) while increasing effector T cells (C24, C27) (Fig. S7E-G), suggesting VLPs drive efficient naïve T cell differentiation into an effector state. Among B cell subsets, activated B cells (C15) differentiated into more *Aicda* and *Ada*-expressing tissue-resident B cells (C18) and proliferating B cells (C17) after Ad5-HA_PR8_-VLP immunization (Fig. S7H-J), indicating that VLP promotes the proliferation and differentiation of lung B cells.

Collectively, these findings demonstrate that Ad5-HA_PR8_-VLP vaccination profoundly reshapes the pulmonary immune landscape by enhancing antigen presentation, metabolic activity, and inflammatory signaling in APCs (particularly DCs and macrophages), promoting novel functional states in NK cells, driving effector T cell differentiation, boosting B cell proliferation and survival via APRIL-mediated communication, and establishing a pro-migratory environment, thereby providing a comprehensive mechanistic basis for its superior immunogenicity.

### Ad5-HA_PR8_-VLP mucosal immunization confers long-term and broad-spectrum protection against influenza A virus

To assess the long-term cross-protective capacity, vaccinated mice were challenged intranasally with homologous (PR8) or heterologous (CA04/H3N2) influenza A viruses at 174 dpi (Fig. 6A). Unvaccinated controls exhibited rapid disease progression, with a mean body weight loss of 26.34% and 100% mortality within 8 days after PR8 challenge (Fig. 6B and C). In contrast, both intranasal (i.n.) and intramuscular (i.m.) Ad5-HA_PR8_-VLP vaccination achieved 100% survival without virus-associated clinical manifestations. Notably, we observed that survival rates in Ad5-HA_PR8_-vaccinated groups showed immunization route-dependent variation: 80% (8/10) for intranasal versus 90% (9/10) for intramuscular administration. Weight loss profiles further distinguished the different immunization strategies, with Ad5-HA_PR8_-VLP nasal immunization-induced body weight loss (maximum loss 11.3%) being significantly lower than that induced by i.m. (17.2%). Similarly, Ad5-HA_PR8_ intranasal-vaccinated mice displayed attenuated weight loss (11.6% maximum) relative to their intramuscular counterparts (19.3%). Additionally, on day 5 post-challenge, Ad5-HA_PR8_-VLP immunized mice demonstrated significantly lower lung viral titers compared to both unvaccinated controls and Ad5-HA_PR8_ immunized groups, with intranasal immunization achieving further reduction in pulmonary viral loads than intramuscular administration (Fig. 6D).

**Figure 6.**
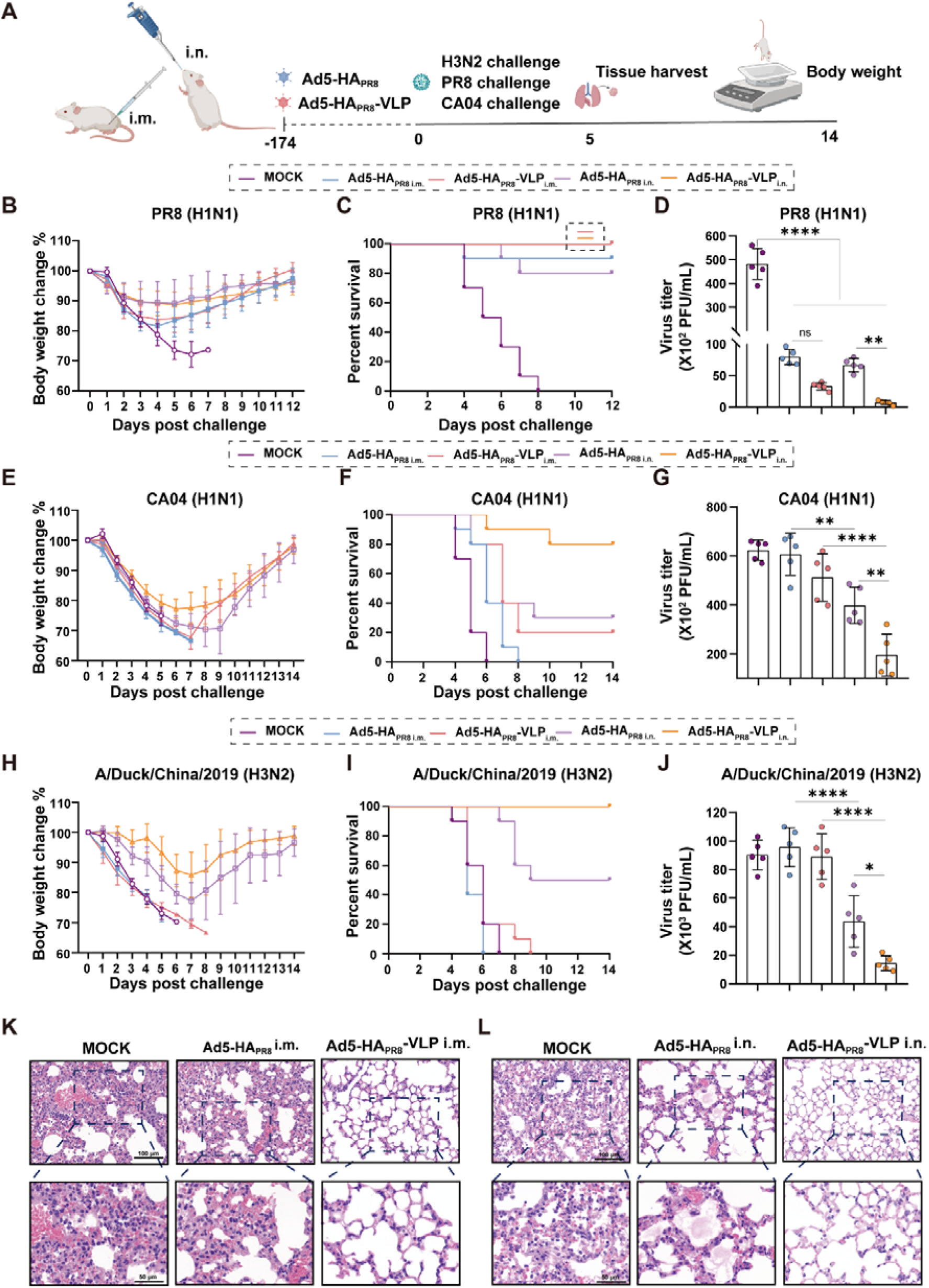
Long-term protective efficacy against lethal challenges with divergent variants of influenza A viruses. (A) Experimental design schematic. BALB/c mice were immunized intranasally or intramuscularly with 10□ TCID□□ Ad5-HA_PR8_-VLP, Ad5-HA_PR8_, or DMEM (vehicle control). Mice were intranasally challenged with 10^4^ PFU PR8 (H1N1), 3×10^4^ PFU CA04 (H1N1), or 10^4^ PFU A/duck/China/Influenza A virus/2019 (H3N2) at 174 dpi, respectively. Lungs (n=5 per group) were collected at 5 days post-challenge for viral titer quantification. Body weight changes and survival rates were monitored for 12–14 days. Body weight changes (B) and survival rates (C) in PR8-challenged cohorts. (D) Lung viral titers at 5 days post-PR8 challenge. Weight loss profiles (E) and Survival (F) following CA04 challenge. (G) Pulmonary viral loads post-CA04 challenge. Weight changes (H) and survival (I) in H3N2-challenged mice. (J) Lung viral titers post-H3N2 challenge. (K) Representative H&E-stained lung sections from PR8-challenged mice immunized intramuscularly with Ad5-HA_PR8_-VLP or Ad5-HA_PR8_. (L) Corresponding sections from intranasally immunized mice. Scale bars: 100 μm. Data are presented as mean ± SD. Survival significance was determined by log-rank (Mantel-Cox) test. Panel D, G and J was analyzed by one-way ANOVA with Tukey’s test. Significance levels: NS (not significant), \**P* < 0.05, \*\**P* < 0.01, \*\*\**P* < 0.005, \*\*\*\**P* < 0.0001.

Following CA04 challenge, unvaccinated controls showed rapid disease progression (24% mean weight loss) with 100% mortality within 6 days post-challenge (Fig. 6E and F). Ad5-HA_PR8_-VLP intranasal immunization conferred 80% survival (8/10) with minimal clinical signs, while intramuscular administration achieved only 20% survival (2/10). In contrast, Ad5-HA_PR8_ intranasally vaccinated mice exhibited 30% survival, and all intramuscularly immunized animals succumbed to infection despite delayed mortality onset. Weight loss profiles paralleled survival trends, intranasal delivery of Ad5-HA_PR8_-VLP limited maximum weight reduction to 22% versus 33% in intramuscularly immunized groups, while Ad5-HA_PR8_ groups showed 29% (i.n.) and 34% (i.m.) losses (Fig. 6E). A similar pattern emerged in A/Duck/China/2019 (H3N2)-challenged cohorts (Fig. 6H and I). Unvaccinated controls reached 100% mortality within 6 dpi. We observed that Ad5-HA_PR8_-VLP intranasal immunization achieved complete survival (100%) with less than 14% weight loss, whereas intramuscular administration showed 100% lethality. Ad5-HA_PR8_ intranasal vaccination showed intermediate efficacy (50% survival, 22% weight loss), while intramuscular delivery again proved ineffective. Virological analysis on day 5 post-challenge revealed enhanced pulmonary protection in Ad5-HA_PR8_-VLP intranasal-immunization groups. Viral titers were 3.2- and 6.2-fold lower than those in unvaccinated controls in CA04 and H3N2 models, respectively (Fig. 6G and J). Notably, intramuscular administration of both vaccines failed to reduce viral loads compared to naïve controls across challenges.

H&E staining of lung tissues from PR8-challenged mice showed distinct immunization-mediated protection patterns (Fig. 6K-L). Mice receiving Ad5-HA_PR8_-VLP immunization maintained normal pulmonary architecture without pathological alterations. In contrast, Ad5-HA_PR8_-vaccinated animals exhibited severe interstitial pneumonia characterized by alveolar septal thickening, perivascular leukocyte infiltration, and bronchiolar luminal exudates. Protective efficacy varied significantly depending on administration route. Intranasal immunization with Ad5-HA_PR8_-VLP resulted in markedly reduced pulmonary damage compared with intramuscular delivery, with fewer inflammatory foci in intranasal immunization groups. Similarly, Ad5-HA_PR8_ intranasal immunization attenuated alveolar hemorrhage severity relative to its intramuscular counterpart, though both routes showed greater pathology than in VLP-vaccinated mice. These results demonstrate that intranasal Ad5-HA_PR8_-VLP immunization confers protection against homologous (PR8) and heterologous (CA04, H3N2) influenza A challenges, as evidenced by reduced viral loads and attenuated pulmonary pathology.

### Ad5-Envp-VLP is a universal platform for pioneering broad-spectrum antiviral vaccine

To investigate the universal applicability of the Ad5-Envp-VLP platform, we evaluated the immunogenicity of two additional vaccine candidates (Ad5-S_JN.1_-VLP and Ad5-RVDG-VLP). Details regarding the construction and characterization of these vaccines are provided in Fig. S8. In the SARS-CoV-2 model, the mucosal immune efficacy of Ad5-S_JN.1_-VLP and Ad5-S_JN.1_ was evaluated. Specifically, mice were intranasally immunized with 10□ TCID_50_ of either Ad5-S_JN.1_ or Ad5-S_JN.1_-VLP, with serum and BALF collected at 14 and 84 dpi (Fig. 7A). S-specific IgA titers in BALF assessed by ELISA showed that the Ad5-S_JN.1_-VLP group induced significantly higher IgA titers than the Ad5-S_JN.1_ group at both day 14 and 84 (Fig. 7B). Pseudovirus neutralization assays revealed broad-spectrum neutralizing activity, with the VLPs formulation generating enhanced antibody titers against the homologous JN.1 strains and heterologous variants including WA1/D614G, B.1.617.2, and BA.2.86 (Fig. 7C). Overall, our results suggest that Ad5-S_JN.1_-VLP induces potent and broad-spectrum antibodies against SARS-CoV-2 variants.

**Figure 7.**
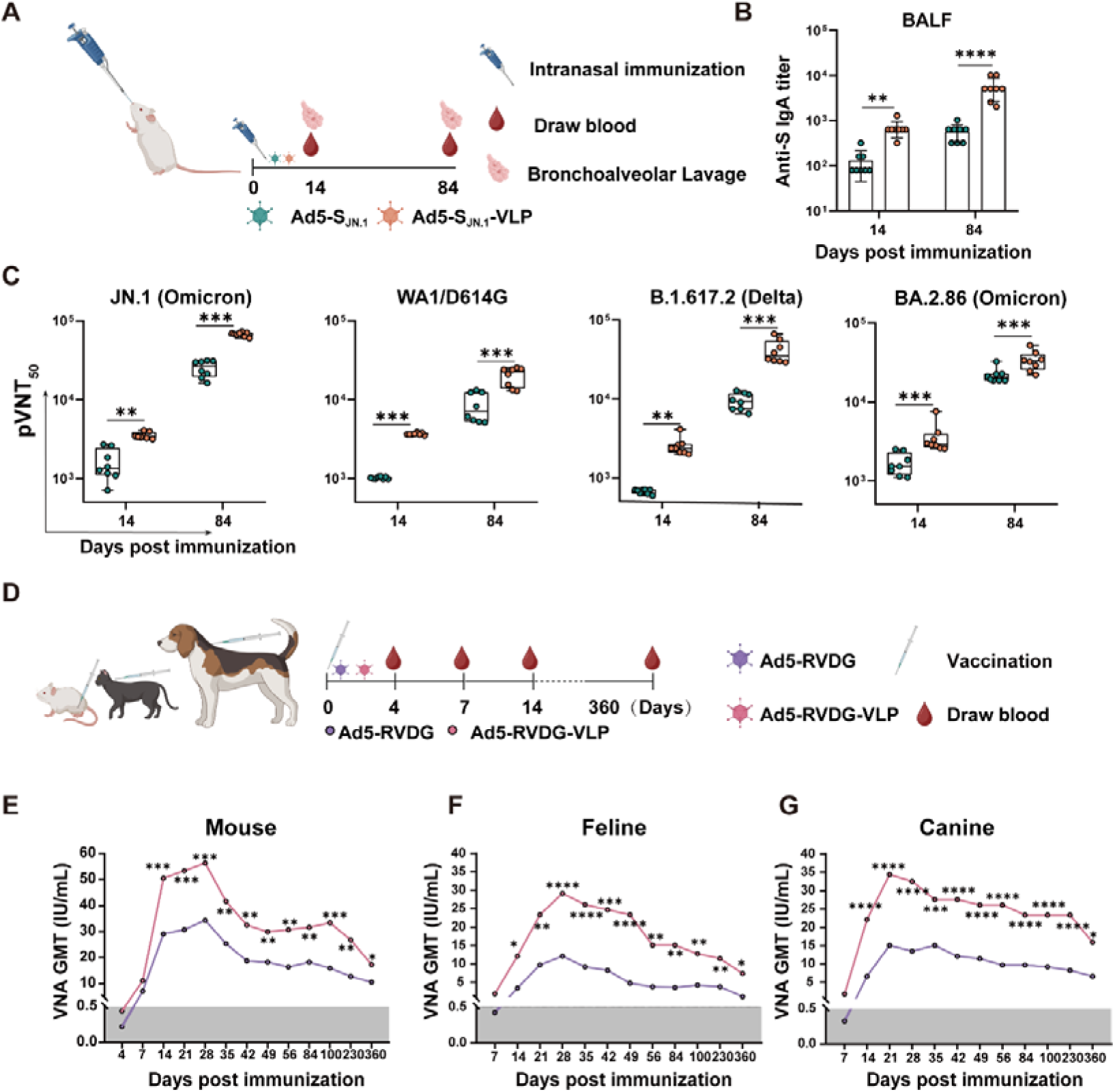
Versatility of the Ad5-Envp-VLP platform. (A) SARS-CoV-2 model: BALB/c mice (n=8 per group) were intranasally immunized with 10□ TCID_50_ Ad5-S_JN.1_-VLP or Ad5-S_JN.1_ (control). Serum and BALF were collected at designated time points. (B) S-specific IgA titers in BALF (n=8 per group) at days 14 and 84. (C) 50% pseudovirus neutralization (pVNT_50_) titers against SARS-CoV-2 variants (JN.1, WA1/D614G, B.1.617.2 and BA.2.86). (D) Rabies virus (RABV) model: ICR mice (n=10 per group) were intramuscularly immunized with 10□ TCID□□ Ad5-RVDG or Ad5-RVDG-VLP Rabies vaccine-naïve beagles and cats (n=5 per group) received subcutaneous injections of 10□ TCID□□ Ad5-RVDG or Ad5-RVDG-VLP in the neck region. Serum was collected for RABV-specific neutralizing antibody quantification. (E) VNA titers in mice over 360 dpi. (F, G) VNA titers in cats and dogs over 360 dpi. Data are presented as mean ± SD. Statistical significance (Panels B, C, E, F and G) was determined by two-way ANOVA with Tukey’s multiple-comparison test. Significance levels: NS (not significant), \**P* < 0.05, \*\**P* < 0.01, \*\*\**P* < 0.005, \*\*\*\**P* < 0.0001.

In the influenza models, we found Ad5-HA_PR8_-VLP induced rapid and durable antibodies by i.m. immunization. Early antibody production is critical for rabies prophylaxis, as rapid neutralization after rabies vaccination determines the protective effect. Currently, the rabies prophylaxis recommended by the World Health Organization still relies on multiple doses of inactivated vaccines (3–5 vaccinations), which show delayed seroconversion (7–10 days after vaccination) and short-term antibody persistence (<1 year). To evaluate the immune effect of the Ad5-Envp-VLP platform in the rabies model, we further constructed Ad5-RVDG-VLP and explored its immune effect on different species (mice, dogs and cats) (Fig. 7D). Our data revealed that Ad5-RVDG-VLP immunization induced rabies-specific neutralizing antibodies (VNA ≥0.5 IU/mL) in 50% of mice by day 4 (versus 2 responder in Ad5-RVDG group), achieving 100% seropositivity by day 7, peaking at day 28 (56.31 versus 34.34 IU/mL in Ad5-RVDG group), and maintaining protective titers (17.28 vs. 10.54 IU/mL) at 360 days (Fig. 7E). Cross-species validation confirmed consistent superiority of the VLP formulation. In feline models, Ad5-RVDG-VLP achieved 100% seroconversion (VNA ≥ 0.5 IU/mL) by day 7, with peak titers of 31.2 IU/mL at day 28 versus 12.2 IU/mL in controls (Fig. 7F). Longitudinal monitoring showed sustained protection in all VLP-immunized cats (GMT >0.5 IU/mL at 360 days), while one control animal declined to non-protective levels (0.38 IU/mL). In canine models, Ad5-RVDG-VLP mirrored the findings observed in other species, achieving 100% seropositivity by day 7 (versus 40% in controls) and maintaining a 2.4-fold higher GMT at the study endpoint (15.92 versus 6.63 IU/mL), despite comparable long-term seroprotection rates between groups (Fig. 7G). In summary, the Ad5-Envp-VLP platform demonstrates robust cross-cutting versatility in vaccine development, characterized by its broad applicability across diverse high-pathogenicity pathogens, immunization modalities, and species.

## DISCUSSION

Vaccination remains the most cost-effective intervention for achieving herd immunity against emerging and re-emerging infectious diseases. In this study, we developed Ad5-Envp-VLP, a recombinant adenovirus platform integrating the EABR strategy to enable spontaneous eVLPs assembly both *in vivo* and *in vitro*. This platform eliminates the need for pathogen-specific assembly scaffolds or laborious post-translational modifications, overcoming a critical limitation of conventional VLP technologies. VLP-based immunization significantly enhances durability of antibody responses against influenza and rabies viruses, resulting in long-term protection against lethal challenges in IAV models. Crucially, mucosal delivery of Ad5-Envp-VLP mediated the recruitment, activation, and transcriptional reprogramming of pulmonary innate immune cells, which triggered superior respiratory sIgA titers and polyfunctional T-cell responses compared to intramuscular administration. This mucosal immunization strategy conferred cross-protection against heterologous influenza strains (Fig. 8). Notably, in SARS-CoV-2 models, the platform further elicited broader and more potent neutralizing antibody titers. In summary, our results demonstrated Ad5-Envp-VLP’s dual competency in antigen delivery and immune reprogramming, offering a rapid-response platform with broad-spectrum efficacy against highly variable envelope viruses.

**Figure 8.**
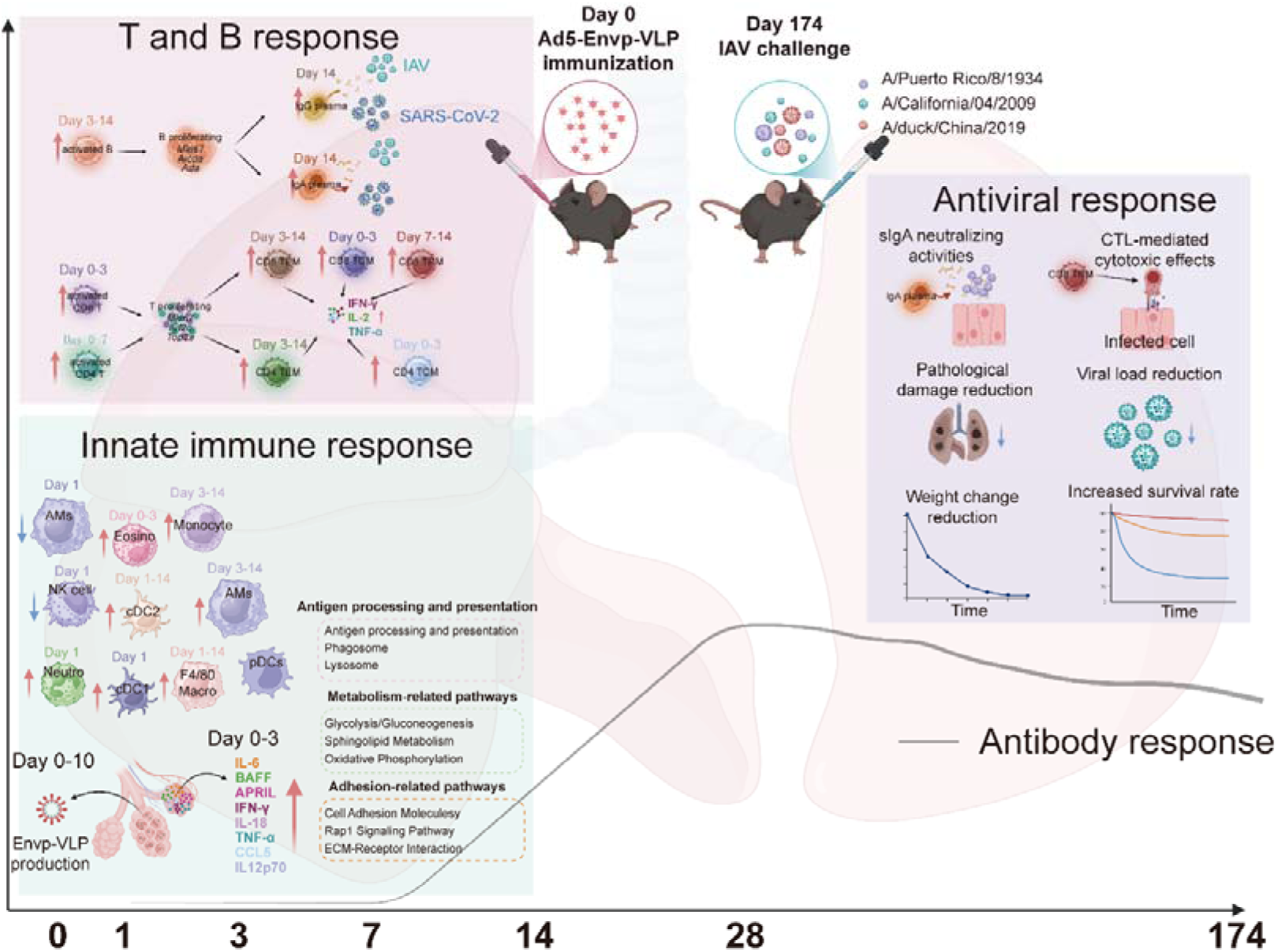
Schematic illustration of Ad5-Envp-VLP induces mucosal immune response *in vivo*. Immune response kinetics and protection following intranasal immunization: Within 0-7 dpi, innate immune cells are highly activated and produce cytokines (including IL-6, BAFF, APRIL, IFN-γ, IL-18, TNF-α, CCL5 and IL-12p70). By 7 dpi, myeloid cells show significant enrichment in pathways related to: antigen processing and presentation (Phagosome, Lysosome), metabolism (Glycolysis/Gluconeogenesis, Oxidative Phosphorylation, Sphingolipid Metabolism), and cell migration/adhesion (Cell Adhesion Molecules, Rap1 Signaling, ECM-Receptor Interaction). Subsequently (3-7 dpi), activated B cell numbers increase significantly, ultimately differentiating into plasma cells secreting pathogen-specific IgA and IgG. Concurrently (0-7 dpi), activated CD4+ and CD8+ T cell numbers increase markedly in the lungs, differentiating into effector memory TEM and central memory (TCM) subsets. By 14 dpi, lung tissue-resident CD8+ TRM are significantly increased. Crucially, Ad5-Envp-VLP confers durable protection against lethal homologous and heterologous influenza virus challenge at 174 dpi, evidenced by reduced lung histopathology, attenuated body weight loss, and decreased lung viral load.

Modern vaccinology combines modular platforms (e.g., mRNA, VLPs, adenoviral vectors) with structural design to amplify immune response by synergizing their complementary strengths, surpassing limitations of single-platform approaches. For instance, rabies VLP/mRNA vaccines co-expressing structural proteins (preG/M/N) enhance germinal center activation compared to monomeric G protein(*8*), and SARS-CoV-2 VLP-encoding mRNA vaccines elicit higher neutralizing antibody titers than membrane-anchored Spike mRNA vaccines(*9*). However, these platforms generally require multiple viral antigens for VLP assembly and exhibit restricted cross-species applicability or antigenic versatility. Although the EABR platform improves antigen density through ESCRT-mediated eVLP assembly(*11*), it has yet to be validated in VLP biogenesis *in vivo*, cross-species translational models, and multi-pathogen efficacy testing. Our Ad5-Envp-VLP system demonstrated universal functionality across human, murine, canine, and feline cells, producing pathogen-matched VLPs with conserved immunogenicity. The evolutionary conservation of ESCRT machinery likely underpins this cross-species efficacy(*32*), as evidenced by superior rabies-neutralizing titers in all tested mammals. The combination of manufacturing scalability and rapid single-dose seroconversion kinetics (<7 days) makes our platform a transformative tool for both epidemic preparedness and veterinary applications.

The rapid development of neutralizing antibodies following vaccination is critical for pathogen clearance in acute infections such as rabies, with early antibody titers serving as validated surrogates for clinical efficacy(*33, 34*). Multivalent antigen presentation enhances BCR crosslinking, amplifying early humoral responses(*2, 3, 35*). Previous studies demonstrated that high-density HIV Env trimers displayed on 60-mer VLPs significantly improve BCR crosslinking efficiency compared to low-valency formats, driving a 4-fold expansion of GC B cells and accelerating neutralizing antibody generation within 6 dpi(*36*). Consistent with this finding, we found that Ad5-Envp-VLP platform elicited stronger antibody responses than soluble Ad5-Envp within 4–7 days in IAV and RABV immunization models, further indicating that antigen display by VLPs induces early antibody production. FCM analysis revealed a 4.39-fold increase in activated B cells in iLNs by day 3, followed by elevated GC B cells and SLPCs by day 7. Transcriptomic profiling showed that VLP immunization upregulated cell cycle regulators (*Cdc6, Cdc45, Plk1, Ccnb1/2, Aurka/b*), suggesting enhanced B cell proliferation, whereas downregulation of IL7R indicated accelerated B cell maturation(*37*). Mechanistically, elevated PI3K activity (driven by increased *Pik3r6* transcription) in VLP-stimulated naïve B cells promotes proliferative responses while paradoxically suppressing CSR and SHM through transcriptional inhibition of *Aicda*(*38*). However, our data showed that Ad5-HA_PR8_-VLP immunization induced a 2.6-fold increase in *Aicda* expression compared to Ad5-HA_PR8_, indicating that VLPs overcome this regulatory constraint to promote CSR and SHM in germinal centers. Notably, adenosine deaminase (ADA), a purine-metabolizing enzyme critical for B cell survival(*39*), was upregulated in Ad5-HA_PR8_-VLP-immunized mice. *ADA* deficiency disrupts dNTP homeostasis, impairing B cell proliferation and function, as evidenced by rescued B cell counts and class-switching capacity in *ADA* gene therapy trials (e.g., OTL-101)(*40*). *ADA* upregulation induced by VLPs suggests metabolic reprogramming to sustain dNTP pools, facilitating GC reactions and antibody diversification. Together, with previous studies, our study demonstrates that multivalent display of VLPs enhances BCR cross-linking, proliferation signaling, and metabolic adaptation, thereby accelerating early antibody production.

Bidirectional innate-adaptive immune crosstalk forms the cornerstone of infection defense, with DCs and Toll-like receptor (TLR) signaling classically driving T cell priming through pathogen sensing(*41, 42*). However, emerging evidence suggests that B cells can directly activate naïve CD4+ T cells independently of DCs, thereby promoting Tfh cell differentiation during RNA phage-derived VLP immunization(*23, 43*). In this study, we demonstrate that immunization route may determines APC engagement dynamics for Ad5-HA_PR8_-VLP. Intramuscular administration induced rapid B cell expansion (7 dpi) with enriched antigen processing and presentation genes, suggesting B cells may dominate early CD4+ T cell priming in this route. Conversely, intranasal delivery elicited robust activation of DCs, AMs, and interstitial macrophages, highlighting mucosal pathway specificity potentially mediated through mucosa-associated lymphoid tissue (MALT) networks.

Pulmonary mucosal immunity is orchestrated by both professional and non-professional APCs(*44*). Previous studies have shown that Ams capture native antigens on their surfaces and serve as effective APCs for naïve B cell activation in the lung(*45*). In this study, scRNA-seq revealed that intranasal Ad5-HA_PR8_-VLP immunization drives macrophage recruitment and upregulates genes linked to antigen endocytosis, processing, and presentation. Macrophage-B cell crosstalk via MIF-CD74/CD44 interactions further enhances B cell activation. Notably, Ad5-HA_PR8_-VLP immunization significantly expanded bronchus-resident memory B cells (BRMs) that established SPP1-CD44 and MIF-CD74/CD44 interactions with AMs. These findings align with those of MacLean *et al*. (*46*), in which BRM-macrophage cooperation during influenza reinfection was reported to coordinate localized immunity. Furthermore, elevated BAFF/APRIL levels at day 7, critical for naïve B cell activation(*47*), IgA class-switching(*48*), and plasma cell survival(*49, 50*)— correlated with macrophage-derived APRIL-TNFRSF17/TNFRSF13B signaling. Our findings align with mechanisms reported by Kawasaki et al. (*51*), where AMs-derived IL-18 promotes CD103+ CD8+ tissue-resident memory T cell (TRM) differentiation to suppress influenza replication. Ad5-HA_PR8_-VLP immunization induced transcriptional upregulation of IL18 in lung macrophages alongside CD8+ TRM expansion, suggesting VLP-driven macrophage activation may enhance T cell lung residency. Consistent with Si et al.(*52*), who identified pulmonary cDC1/cDC2 subsets as critical mediators of intranasal nanofiber-induced CD8+ T cell priming prior to lymph node trafficking. Our study revealed that Ad5-HA_PR8_-VLP immunization enhances DC-dependent antigen cross-presentation. Transcriptional profiling of DCs from Ad5-HA_PR8_-VLP immunized mice showed significant upregulation of MHC class I antigen presentation genes (*PDIA3, KLRD1, HSPA5, HSPA8, HSP90AB1*), indicating augmented DC-mediated antigen processing. This finding aligns with the established role of DC in cross-presentation and supports the broader applicability of mucosal VLP platforms in eliciting cytotoxic T cell responses. Notably, we observed elevated *IL18* transcription in DCs post-immunization. While IL-18 is known to recruit pDCs via IL-18R signaling during early innate responses(*53, 54*), FCM revealed no significant changes in pDC proportions. However, scRNA-seq identified an expanded pDC population with upregulated genes linked to phagocytosis, oxidative phosphorylation (OXPHOS), and antigen presentation. These findings mirror observations by Wu et al.(*55*), wherein TLR9-activated pDCs undergo metabolic reprogramming (FAO/OXPHOS upregulation) to sustain IFN-α/β production. The discordance between FCM and transcriptomic data may reflect enhanced metabolic activity rather than numerical expansion, suggesting VLPs potentiate pDC functionality through energy metabolism remodeling. Emerging evidence underscores neutrophils’ ability to act as non-canonical APCs during direct interactions with lymphocytes(*56–58*). IFN-γ and GM-CSF stimulation induces MHC II and co-stimulatory molecule expression (CD80/86, CD83) in neutrophils(*56, 59*), enabling APC-like functionality—a property also observed in freshly isolated human neutrophils(*60*). In Ad5-HA_PR8_-VLP intranasal immunization model, pulmonary neutrophils exhibited upregulated phagosome-related (*Mrc1*, *Cybb*) and MHC II pathway genes (*H2-Ab1*, *Cd74*, *H2-Eb1*), indicative of enhanced antigen-processing capacity. Furthermore, neutrophil-derived FOXO1 transfer to naïve T cells—a mechanism linked to FOXP3+CD4+ Treg differentiation with IL-10, VEGF and IL-17 secretion(*61*) — was corroborated by elevated FoxO signaling activity and expanded Treg populations in immunized mice (Fig. S7F), suggesting a neutrophil-driven immune regulatory axis. Concurrently, we identified a novel NK cell subset expressing MHC II-associated genes (*Cd74, H2-Eb1, H2-Ab1*) after Ad5-HA_PR8_-VLP immunization. While NK cells are traditionally recognized for cytotoxicity and cytokine-mediated antiviral responses(*54*), the dual phenotypic traits of this subset imply functional diversification, potentially bridging innate pathogen sensing with adaptive immune modulation.

Emerging evidence have established pulmonary mucosal immunity as a frontline defense against antigenically plastic respiratory pathogens, including influenza and SARS-CoV-2(*62, 63*). Tutykhina et al(*64*) demonstrated that an Ad5-vectored vaccine expressing influenza M2 and NP epitopes (Ad5-tet-M2NP) conferred durable protection against five divergent influenza subtypes, emphasizing the value of conserved antigenic targets. Similarly, HMNF nanoparticles displaying multivalent influenza epitopes achieved complete heterosubtypic protection by inducing broad humoral and mucosal responses(*65*). In line with these advances, our findings demonstrated that intranasal Ad5-HA_PR8_-VLP, rather than soluble Ad5-HA_PR8_, elicited cross-protection against heterologous influenza strains, likely via enhanced lung-resident B cell and antigen-specific CD8+ T cell responses. The critical role of mucosal priming is further underscored by comparative different vaccine platforms. Live attenuated influenza virus (LAIV) vectors expressing SARS-CoV-2 RBD outperformed intramuscular mRNA vaccines in reducing viral replication and pathology(*66*). Analogously, our data revealed that intramuscular Ad5-HA_PR8_-VLP immunization conferred homologous but not heterologous protection, with elevated early-phase lung viral loads in heterosubtypic challenges. In sharp contrast, intranasal Ad5-HA_PR8_-VLP achieved cross-protection, highlighting the necessity of mucosal priming for frontline defense. The failure of intramuscular Ad5-HA_PR8_-VLP to control heterologous infection despite systemic immunity further underscores the compartmentalized nature of respiratory defense, where mucosal responses uniquely intercept respiratory pathogens at portals of entry.

In conclusion, we establish the Ad5-Envp-VLP platform that synergizes adenoviral mucosal tropism with multivalent antigen presentation through VLP assembly. This engineered system enables *in situ* pulmonary VLP biogenesis, triggering a coordinated innate-adaptive immune cascade characterized by (1) mucosal sIgA production, (2) TRM establishment, and (3) rapid neutralizing antibody responses—critical determinants of cross-protection against heterologous influenza strains. Notably, intramuscular administration demonstrates early seroconversion (3–7 dpi), a vital feature for post-exposure prophylaxis in rabies and other acute lethal infections. To address pre-existing Ad5 immunity — a key translational challenge — capsid engineering strategies, including hypervariable region replacement in Hexon or fiber redesign, enable evasion of neutralizing antibodies(*67*). Leveraging rare-serotype adenoviruses (e.g., Ad26, Ad48) further enhances universal applicability by adapting to regional seroprevalence patterns(*68*). Optimized iterations of this platform hold promise for accelerating vaccines against emerging respiratory pathogens, such as avian influenza variants or SARS-CoV-2 sublineages, shifting global health strategies from reactive containment to proactive interception through mucosal-primed, cross-protective immunity.

## MATERIALS AND METHODS

### Cells and viruses

HEK293 cells (ATCC CRL-1573), HEK293T cells (ATCC CRL-3216), MDCK cells (ATCC, CCL-34), CRFK cells (ATCC, CCL-94) and BSR cells, a cloned cell line derived from BHK-21 cell (ATCC, CCL-10) were maintained in our laboratory. The HEK293-ACE2-OE cell line was obtained from Biodragon Inc. (Cat. No. BDAA0039). These cell lines were cultured in Dulbecco’s modified Eagle’s medium (DMEM, BioChannel Biotechnology Co., Ltd) containing 10% fetal bovine serum (FBS, AusGeneX, Australia) and 1% penicillin/streptomycin (BioChannel Biotechnology Co., Ltd). Kop293 cells were cultured in Kop293 cell culture (KAIRUI BIOTECH Co., Ltd.). Influenza virus A/Puerto Rico/8/1934 (PR8, H1N1) was kindly donated by Dr. Hongbo Zhou (Huazhong Agricultural University, Wuhan, China) and propagated in the allantoic cavity of SPF eggs. Influenza virus A/California/04/2009 (CA04, H1N1) was constructed using reverse genetics and adapted to mice through ten generations of serial passage. Influenza A virus A/duck/China/2019 (H3N2) (kindly provided by Dr. Guoyuan Wen, Hubei Academy of Agricultural Sciences) was adapted through ten generations of serial passages. Replication-deficient adenovirus serotype 5 vector expressing firefly luciferase and eGFP (Ad5-Luci and Ad5-eGFP) and a rabies challenge virus strain CVS-11were preserved in our laboratory.

### Animals and ethics statement

Female BALB/c, C57BL/6, and ICR mice (6–7 weeks old) were procured from the Hubei Provincial Center for Disease Control and Prevention and maintained under specific pathogen-free (SPF) conditions at Huazhong Agricultural University’s Laboratory Animal Center. Influenza virus and rabies virus challenges were conducted in the university’s Animal Biosafety Level 2 (ABSL-2) facility, with humane endpoints defined as ≥30% body weight loss, severe lethargy, respiratory distress, paralysis, or failure to eat/drink. Cats were sourced from Jiaxiang Huarong Breeding Cooperative, and 3–5-month-old beagles were obtained from Yizhicheng Biological Technology Co., Ltd. (Yingcheng city, Hubei, China). All experimental protocols were approved by the Scientific Ethics Committee of Huazhong Agricultural University (Approval IDs: HZAUMO-2024-0033, HZAUCA-2024-0007, HZAUDO-2024-0003). Animals meeting endpoint criteria were euthanized via CO□ asphyxiation followed by cervical dislocation.

### Structural design of chimeric fusion proteins

The engineered chimeric protein was constructed by fusing the EABR domain (residues 160–217) of human CEP55 protein with the endocytosis prevention motif (EPM) derived from the cytoplasmic tail of murine Fcγ receptor FcgRII-B1, linked via a flexible 2×GGGS spacer to ensure spatial separation between functional modules(*11*). To enable membrane anchoring and cellular trafficking capabilities, the following viral envelope glycoproteins were independently fused to the N-terminus of the resulting fusion protein: (1) hemagglutinin (HA; residues 1–552) from the influenza A virus PR8 strain (H1N1); (2) glycoprotein (RVG; residues 1–480) of the rabies virus SAD-L16 strain; and (3) spike protein (S; residues 1–1232) of the SARS-CoV-2 JN.1 variant (Omicron sublineage). These constructs were designated HA-EABR, RVG-EABR, and S-EABR, respectively.

### Construction of recombinant adenoviruses

Recombinant adenoviruses were generated using the AdMax™ adenoviral vector system (Microbix Biosystems Inc.), which employs a Cre/loxp-mediated homologous recombination strategy between two plasmids: the backbone plasmid pBHGcre/loxp and the shuttle plasmid pDC315. The chimeric genes encoding HA-EABR, RVG-EABR and S-EABR were cloned into the multiple cloning sites (MCS) of pDC315. These constructs were driven by the human cytomegalovirus (HCMV) immediate-early promoter to generate pDC315-HA-EABR, pDC315-RVG-EABR, and pDC315-S-EABR. Correspondingly, full-length viral membrane proteins (HA, RVG, and S) were similarly cloned into the pDC315, yielding pDC315-HA, pDC315-RVG, and pDC315-S. To enhance exogenous gene expression, the HCMV-driven expression cassettes from the pDC315-derived plasmids were subcloned into the E3 region of pBHGcre/loxp via restriction-ligation, following established protocols(*20*), to generate pBHGcre/loxp-HA, pBHGcre/loxp-HA-EABR, pBHGcre/loxp-RVG and pBHGcre/loxp-RVG-EABR. Pairs of backbone and shuttle plasmids (Table. S1) were co-transfected into HEK293 cells using jetPRIME® (Polyplus). At 8 days post-transfection, supernatants containing primary viral particles were harvested and used to infect fresh HEK293 monolayers. Cytopathic effects (CPE), characterized by cell rounding and detachment, were monitored daily. Six replication-incompetent recombinant adenoviruses were successfully rescued and designated as Ad5-HA_PR8_, Ad5-HA_PR8_-VLP, Ad5-RVDG, Ad5-RVDG-VLP, Ad5-S_JN.1_ and Ad5-S_JN.1_-VLP.

### Growth kinetics analysis of recombinant adenoviruses in HEK293 cells

HEK293 cells were seeded into 12-well plates and cultured until reaching 90% confluency. Cells were infected with recombinant adenoviruses at a multiplicity of infection (MOI) of 0.1 in serum-free DMEM. Maintenance medium containing 2% FBS was replenished 1 hour post-infection (hpi). Viral supernatants were harvested at 12-h intervals (0–96 hpi) for titration. Supernatants were serially diluted (10-fold gradients in DMEM supplemented with 2% FBS) and inoculated into 96-well plates containing HEK293 monolayers (100 µL/well, n=8 replicates per dilution). At 72 hpi, supernatants were discarded, and cells were gently washed twice with phosphate-buffered saline (PBS). Infected cells were fixed with 4% paraformaldehyde for 30 min at room temperature (RT), followed by permeabilization in PBS containing 0.1% Triton X-100 for 15 min. After three PBS washes, cells were blocked with 2% BSA in PBS for 2 h at RT and incubated with an anti-adenovirus hexon monoclonal antibody (Clone 3G0) for 2 h at 37°C. Following two PBS washes, cells were incubated with Alexa Fluor™ 488-conjugated goat anti-mouse IgG (H+L) cross-adsorbed secondary antibody for 45 min and washed twice with PBS. Antigen-positive foci were visualized using an Olympus IX51 fluorescence microscope. Viral titers were calculated as 50% tissue culture infective dose per milliliter (TCID_50_/mL) via the Reed-Muench method.

### Western blotting

Western blotting was performed to analyze exogenous gene expression at 48 hpi. Cell supernatants and lysates from adenovirus-infected cells were collected separately, mixed with SDS loading buffer at a 4:1 (v/v) ratio, and boiled at 95°C for 10 min. Proteins were resolved on 10% SDS–PAGE gels and electrophoretically transferred to 0.45 μm PVDF membranes (Bio-Rad) using a semi-dry transfer system. Membranes were blocked with 5% (w/v) nonfat dry milk in Tris-buffered saline with 0.1% Tween-20 (TBST) for 3 h at RT, followed by overnight incubation at 4°C with primary antibodies diluted in blocking buffer. After three 10-min washes with TBST, membranes were incubated with HRP-conjugated secondary antibodies (1:5,000 dilution) for 1 h at 25°C. Protein signals were developed using the BeyoECL Star chemiluminescence kit (Cat. No. P0018A, Beyotime) and quantified with an Amersham Imager 600 system (GE Healthcare).

### Purification of recombinant adenovirus and virus-like particles

Recombinant adenoviruses and VLPs were produced by infecting HEK293-derived Kop293 cells at a MOI of 2, followed by incubation for 72 h at 37°C with 5% CO_2_. Viral supernatants were clarified via centrifugation (5,000 × g, 30 min, 4°C) to remove cellular debris, then co-purified through iodixanol density gradient ultracentrifugation using an Optima™ TLX ultracentrifuge (Beckman Coulter) with a TLA100.3 rotor (120,000 × g, 2 h). Fractions enriched with adenovirus (density ∼1.34 g/mL) or VLPs (1.18–1.25 g/mL) were pooled and dialyzed against 50 mM NaPO_4_, 65 mM NaCl, 0.005% Tween-80 (pH 6.0) at 4°C for 24 h. For *in vivo* applications and transmission electron microscopy (TEM), VLP preparations underwent additional size-exclusion chromatography (SEC) refinement using a Superose™ 6 Increase 10/300 GL column (Cytiva) equilibrated with PBS (pH 7.4) at a flow rate of 0.5 mL/min.

### Transmission Electron Microscopy (TEM) analysis of VLPs morphology

300-mesh copper grids coated with Formvar/carbon (Electron Microscopy Sciences) were glow-discharge for 30 s to enhance hydrophilicity. Excess liquid was removed, and samples were air-dried prior to imaging. Grids were imaged on transmission electron microscope (Hitachi, Japan) operated at 80.0 kV.

### Mouse immunization and challenge test

In the IAV model, BALB/c mice (n=10/group) were immunized via two routes: (1) intramuscular (i.m.) injection with 10^7^ TCID_50_ of Ad5-HA_PR8_, Ad5-HA_PR8_-VLP or DMEM in 100 μL, and (2) intranasal inoculation under isoflurane anesthesia with an equivalent dose (10^7^ TCID_50_ in 40 μL) of the same vaccines. Viral challenge was performed 174 days post-primary immunization via intranasal administration of 10^4^ PFU PR8 (H1N1 strain), 3×10^4^ PFU CA04 (H1N1 strain), or 10^4^ PFU H3N2 virus. Body weight was monitored daily for 12–14 consecutive days post-challenge, with humane euthanasia implemented for mice exhibiting >30% weight loss to meet ethical endpoints.

In the rabies virus model, ICR mice (n=10/group) were immunized via hind-limb intramuscular injection with 10^7^ TCID_50_/100 μL of Ad5-RVDG, Ad5-RVDG-VLP, or DMEM control. Besides, 3–4-month-old Beagle dogs and Chinese domestic cats (n=5/group) were received cervical subcutaneous injections of 10^8^ TCID_50_ Ad5-RVDG or Ad5-RVDG-VLP in 1 mL volumes. Weekly cephalic vein blood samples were collected for serum separation, with aliquots stored at −80°C until analysis.

In the SARS-CoV-2 model, BALB/c mice (n=8/group) were intranasally immunized with 10^7^ TCID_50_ Ad5-S_JN.1_ or Ad5-S_JN.1_-VLP in 40 μL, with DMEM serving as control. Retro-orbital blood sampling was performed periodically to obtain serum, which was immediately frozen at −80°C for subsequent immunological assessments.

To compare the immunogenicity of purified VLPs with soluble proteins, 6-week-old female mice were intramuscularly immunized with DMEM, 10 μg AS03-adjuvanted Envp or Envp-VLP. Mouse strain and group sizes were consistent with those used in the three aforementioned pathogen models (H1N1 influenza, rabies virus, and SARS-CoV-2).

### Hemagglutination inhibition (HI) assay

The HI assay was performed according to standardized WHO protocols with minor modifications. Briefly, serum samples were heat-inactivated at 56°C for 30 min. In parallel, influenza A/PR/8/34 (H1N1) virus was titrated in V-bottom 96-well plates to determine 4 hemagglutinating units (4 HAU), defined as the highest viral dilution inducing complete hemagglutination using 1% (v/v) chicken erythrocytes (SenBeJia Biological Technology Co., Ltd.). Serial two-fold serum dilutions (1:8 to 1:1024) were prepared in duplicate using PBS. Each diluted serum sample was mixed with 25 μL of 4 HAU virus suspension per well and incubated at RT for 45 min to facilitate antibody-antigen binding. Finally, 50 μL of 1% chicken erythrocyte suspension was added to each well, followed by a 45-min incubation at 4°C to stabilize hemagglutination patterns.

### Flow cytometry

To identify total GC B cells, Tfh, short-lived plasmacytes and activated B cell, freshly isolated cells (1 × 10^6^ cells/100 μl) from the inguinal lymph nodes (iLNs) were washed 1× with PBS (350 × g, 7 min, 4 °C). Cells were washed 3× followed by incubation with Fc Block (1 μL/test, Cat. No. 14-0161-85, eBioscience™) for 15 min at 4 °C. Samples were incubated for an additional 30 min upon addition of the surface cocktail containing the following anti-mouse antibodies: CD45R (0.5µg/test, clone RA3-6B2, eBioscience™), CD3 (0.5 µg/test, clone 17A2, eBioscience™), CD4 (0.5 µg/test, clone GK1.5, eBioscience™), CD185 (0.25 µg/test, clone L138D7, BioLegend), PD-1 (0.8 µg/test, clone RMP1-30, BioLegend), GL7 (0.5 µg/test, clone GL7, eBioscience™), CD95 (0.25 µg/test, clone SA367H8, BioLegend), CD8a (0.25 µg/test, clone 53-6.7, eBioscience™), CD138 (0.5 µg/test, clone 281-2, BioLegend), CD44 (0.25 µg/test, clone IM7, eBioscience™), CD69 (0.25 µg/test, clone H1.2F3, eBioscience™). For quantitative analysis of HA-specific GC B cells and memory B cells (MBCs), the following protocol was applied: recombinant HA-His protein was labeled with biotin using a Biotinylation Kit (Cat. No. G-MM-IGT, Genemore). Then, single-cell suspensions were incubated with 5 μg/mL biotinylated His-HA (His-HA-Biotin) at 4°C in the dark for 30 min, followed by PBS washing. Cells were subsequently stained with PE-conjugated streptavidin (Cat. No. 405203, BioLegend) under identical conditions (4°C, 30 min) prior to FCM analysis.

Following intranasal immunization with Ad5-HA_PR8_ or Ad5-HA_PR8_-VLP, lungs were harvested at designated timepoints, with lung tissue undergoing mechanical dissociation using the gentleMACS system (Lung_01_02 program), followed by enzymatic digestion using Lung Dissociation Kit, Mouse. (Cat. No. 130-095-927, Miltenyi Biotec™) for 30 min. Tissues were further dissociated (gentleMACS Lung_02_01 program), filtered through a 40-μm strainer, and RBC-depleted using ACK lysis buffer (Cat. No. BL503B, Biosharp™). Cells were washed 3 times followed by incubation with Fc Block for 15 min at 4°C. Samples were incubated for an additional 30 min upon addition of the surface cocktail containing the following anti-mouse antibodies: CD45 (0.125 µg/test, clone 30-F11, eBioscience™), CD3 (0.3 µg /test, clone 17A2, eBioscience™), CD19 (0.3 µg/test, clone 1D3, eBioscience™), CD317 (0.25 µg/test, clone eBio927, eBioscience™), CD8a (0.25 µg/test, clone 53-6.7, eBioscience™), CD11b (0.25 µg/test, clone M1/70, eBioscience™), CD11c (0.5 µg/test, clone N418, eBioscience™), CD86 (0.25 µg/test, clone GL-1, eBioscience™), MHC II (0.025 µg/test, clone M5/114.15.2, BioLegend), Ly6C (0.125 µg/test, clone HK1.4, eBioscience™), Ly6G (0.3 µg/test, clone 1A8, eBioscience™), CD103 (0.5 µg/test, clone 2E7, eBioscience™), F4/80 (0.25 µg/test, clone BM8, eBioscience™), CD169 (0.25 µg/test, clone 3D6.112, BioLegend), Siglec F (0.16 µg/test, clone 1RNM44N, eBioscience™), NK1.1 (0.5 µg/test, clone PK136, eBioscience™) for 30 min at 4°C.

To identify total activated B/T cell, memory T cell, freshly single cells from lungs were then stained with a surface antibody cocktail: CD45 (0.125 µg/test, clone 30-F11, eBioscience™), CD45R (0.5 µg/test, clone RA3-6B2, eBioscience™), CD3 (0.5 µg/test, clone 17A2, BioLegend), CD4 (0.125 µg/test, clone GK1.5, eBioscience™), CD8a (0.25 µg/test, clone 53-6.7, eBioscience™), CD44 (0.25 µg/test, clone IM7, eBioscience™), CD62L (0.25 µg/test, clone MEL-14, eBioscience™), CD138 (0.5 µg/test, clone 281-2, BioLegend), CD69 (0.5 µg/test, clone H1.2F3, eBioscience™), CD103 (0.5 µg/test, clone 2E7, BioLegend) for 30 min at 4°C in the dark.

In the above assay, live cells were identified using 7-AAD Viability Staining Solution (Cat. No. 00-6993-50, eBioscience™). Cells were collected using Cytek Aurora/NL and data were analyzed by FlowJo software V_10.

### ScRNA-seq experimental method

Following intranasal immunization with Ad5-HA_PR8_ or Ad5-HA_PR8_-VLP, lungs were harvested at 7 dpi. Lung tissues were mechanically dissociated using the gentleMACS™ Octo Dissociator (Lung_01_02 program; Miltenyi Biotec), followed by enzymatic digestion with a Lung Dissociation Kit for mice (Cat. No. 130-095-927, Miltenyi Biotec) for 30 min at 37°C. Tissues were further dissociated (gentleMACS™ Lung_02_01 program), filtered through a 40-μm nylon strainer, and subjected to red blood cell (RBC) lysis using ACK lysis buffer (Cat. No. BL503B, Biosharp). Single-cell suspensions were resuspended in ice-cold MACS^®^ buffer (PBS + 0.5% BSA + 2 mM EDTA) and incubated with CD45 MicroBeads (Cat. No. 130-110-618, Miltenyi Biotec) for 15 min at 4°C. After washing with 2 mL MACS® buffer and centrifugation (300 ×g, 5 min), cells were filtered through a 30-μm pre-separation filter and magnetically sorted using a pre-wetted MS column on a MACS® Separator (Miltenyi Biotec). Purified cells were resuspended in 1 mL RPMI 1640 medium (Cat. No. 10-040-CVR, Corning) supplemented with 0.04% (w/v) BSA. Single-cell concentration and viability (>85% viability threshold) were quantified using the LUNA-FL™ automated cell counter (Logos Biosystems, South Korea). Cell suspensions were adjusted to 700–1,200 cells/μL for downstream processing. Libraries were prepared using the 10× Genomics Chromium Next GEM Single Cell 3’ Reagent Kit v3.1 (Cat. No. 1000268) per manufacturer protocols and sequenced on the BGI DNBSEQ-T7 platform (PE100 mode).

### ScRNA-seq data processing

FASTQ files were aligned to the GRCm39 mouse reference genome using Cell Ranger software (v9.0.0; 10× Genomics), and unique molecular identifier (UMI) counts were quantified to generate a cellular barcode expression matrix for downstream analysis with the Seurat package (v4.0.0). Low-quality cells and putative doublets were removed through a five-tier quality control protocol (genes <200, UMIs <1,000, log□□(genes/UMI) <0.7, mitochondrial UMI proportion >10%, hemoglobin UMI proportion >5%) supplemented by DoubletFinder (v2.0.3) prediction. UMI counts were log-normalized (scale factor = 10,000) and the top 2,000 highly variable genes (HVGs) were identified using the FindVariableFeatures function (selection.method = ‘vst’). Dimensionality reduction via principal component analysis (PCA, npcs = 50) enabled graph-based clustering (resolution = 0.5), followed by Uniform Manifold Approximation and Projection (UMAP, dims = 1:20) for 2D visualization. Cluster-specific marker genes were identified using FindAllMarkers (test.use = ‘presto’), while differentially expressed genes (DEGs) were defined by thresholds of |log_2_ (fold change)| >0.58 and adjusted p-value <0.05 (Benjamini-Hochberg correction). Gene Ontology (GO) and Kyoto Encyclopedia of Genes and Genomes (KEGG) pathway enrichment analyses were performed via clusterProfiler (v4.0.5) using hypergeometric testing (pAdjustMethod = ‘BH’). Single-cell sequencing and bioinformatic analyses were conducted by OE Biotech Co., Ltd. (Shanghai, China).

### Cell–cell interaction analysis

Cell-cell communication analysis was performed using the CellChat R package (v2.1.2). First, the normalized expression matrix was imported to create a CellChat object via the createCellChat function. Data preprocessing was then executed through sequential operations: identifyOverExpressedGenes (threshold: p < 0.05), identifyOverExpressedInteractions (species = ‘mouse’), and projectData (reduction.type = “PCA”), all using default parameters. Cell-cell communication probabilities were computed using computeCommunProb (type = “trimean”, distance.method = “cosine”), followed by interaction filtering with filterCommunication (min.cells = 10) to exclude low-confidence signals. Pathway-level communication networks were subsequently resolved via computeCommunProbPathway.

Finally, the integrated cell communication network was aggregated using aggregateNet (remove.isolate = TRUE, threshold.weight = 0.1).

### ELISpot assay

HA-specific antibody-secreting cells (ASCs) were quantified using Multiscreen® HTS ELISpot plates (MilliporeSigma, MA, USA). Plates were coated with 2 μg/mL purified influenza HA-His protein overnight at 4°C, washed three times with PBS, and blocked with RPMI 1640 medium supplemented with 10% FBS for 2 h at 37°C. Single-cell suspensions derived from lung tissues or iLNs were added to the plates and incubated for 24 h under standard culture conditions (37°C, 5% CO□). Following incubation, cells were lysed with ice-cold deionized water for 10 min. Subsequent steps included sequential incubations with: biotinylated anti-mouse IgA/IgG (1:10,000 dilution; Bethyl Laboratories, TX, USA) for 2 h at RT; streptavidin-alkaline phosphatase conjugate (1:1,000 dilution; Mabtech, Sweden) for 1 h at RT; BCIP/NBT Plus substrate (Thermo Fisher Scientific) for 20 min in the dark. Spot-forming units (SFUs) were enumerated using an AID ELISpot Reader (Autoimmun Diagnostika GmbH, Germany). Data normalization was performed by expressing ASC counts per 10□ viable cells, as determined by trypan blue exclusion assay.

### ELISA assays

Antigen-specific IgG, IgA, and IgM antibody titers against influenza HA or SARS-CoV-2 spike (S) proteins were quantified by enzyme-linked immunosorbent assay (ELISA). High-binding 96-well plates (Corning) were coated overnight at 4°C with 2 μg/mL recombinant HA or S protein in carbonate-bicarbonate buffer (pH 9.6), followed by blocking with 5% non-fat milk in PBS containing 0.05% Tween-20 (PBS-T) for 1 h at 37°C. Serially diluted mouse serum or bronchoalveolar lavage fluid (BALF) samples were incubated for 2 h at 37°C. After three washes with PBS-T, plates were incubated with horseradish peroxidase (HRP)-conjugated anti-mouse IgG (1:10,000), IgA (1:5,000), or IgM (1:8,000) (Biodragon) for 1 h at 37°C. Reactions were developed using 3,3’,5,5’-tetramethylbenzidine (TMB) substrate (Solarbio, PR1200; 100 μL/well) for 15 min in the dark, terminated with 100 μL of 2 M H□SO□, and absorbance was measured at 450 nm using a Spark® microplate reader (Tecan). Endpoint titers were defined as the highest serum dilution yielding an optical density (OD) value ≥1.8-fold higher than the negative control (naïve mouse serum).

For HA quantification in BALF from mice intranasally immunized with Ad5-eGFP, Ad5-HA_PR8_, or Ad5-HA_PR8_-VLP, a PR8 HA-specific ELISA kit (H1N1 A/Puerto Rico/8/1934 HA; Sino Biological, KIT11684) was employed. Samples were preprocessed via dilution to optimize detection ranges, per manufacturer instructions. Briefly, PR8 HA-specific monoclonal antibodies pre-coated on plates captured antigens from standards or BALF samples during a 2-h incubation at 25°C. After three PBS-T washes, plates were incubated with HRP-conjugated detection antibody (1:1,000 dilution, 1 h, 25°C), washed again, and developed with TMB substrate (Beyotime, P0209; 100 μL/well, 15 min). Reactions were stopped with 2 M H□SO□, and absorbance was measured at 450 nm using a Spark® microplate reader (Tecan). A linear standard curve (R² >0.99) was generated to calculate HA protein concentrations.

The concentrations of cytokines (including IL-6, BAFF, APRIL, IFN-γ, IL-18, TNF-α, IL-12p70 and CCL5) in lung homogenates from intranasally immunized mice were measured using commercial ELISA kits (Hangzhou Lianke Biotechnology Corp., Ltd., Hangzhou, China).

### Ex vivo stimulation of HA-specific T cells

Lungs and spleens were harvested from mice at specified time points. Single-cell suspensions were isolated via Percoll™ gradient centrifugation (70–40% interface; Cytiva, Cat. No. 17089102) following tissue dissociation. Either lung-derived or spleen-derived single-cell suspensions were plated in complete RPMI-1640 medium at 2–3 × 10□ cells per well in 12-well plates and stimulated for 18 h at 37°C (5% CO□) with: negative control: RPMI-1640 supplemented with 10% FBS (Gibco); antigen stimulation: 20 μg/mL homologous HA protein. Protein transport inhibitors GolgiStop™ and GolgiPlug™ (BD Biosciences) were added during the final 6 h of stimulation. Cells were washed twice with PBS and stained for viability using the Zombie Aqua™ Fixable Viability Kit (Cat. No. 423101, BioLegend), followed by surface staining with anti-CD3 (clone 17A2, eBioscience™), anti-CD4 (clone GK1.5, eBioscience™), anti-CD8a (clone 53-6.7, eBioscience™), anti-B220 (clone RA3-6B2, eBioscience™), anti-CD44 (clone IM7, eBioscience™), and anti-CD62L (clone MEL-14, BioLegend) antibodies. Cells were fixed with intracellular (IC) fixation buffer (Cat. No. 00-5223-56, eBioscience™) for 30 min at RT, followed by permeabilization using intracellular staining permeabilization wash buffer (Cat. No. 00-8333-56, eBioscience™). Intracellular cytokines were stained with anti-IL-2 (clone JES6-5H4, eBioscience™), anti-TNF-α (clone MP6-XT22, BioLegend), and anti-IFN-γ (clone XMG1.2, eBioscience™) antibodies (30 min, 4°C). Samples were acquired on a Cytek Aurora/NL spectral flow cytometer and analyzed using FlowJo™ software (v10.9).

### Immunofluorescence staining

For immunofluorescence staining, tissues were embedded in Optimal Cutting Temperature (OCT) compound (Cat. No. 4583, SAKURA), flash-frozen in liquid nitrogen, and sectioned into 20-μm-thick slices using a Leica CM1950 cryostat (Leica Biosystems, Heerbrugg, Switzerland). Sections were blocked with 10% (v/v) goat serum in PBS for 2 h at RT. Germinal centers in iLNs were stained using a cocktail of the following antibodies (all from BioLegend, unless noted): anti-B220 (clone RA3-6B2, 10 μg/mL), anti-IgG (clone Poly4053, 4 μg/mL), and anti-GL7 (clone GL7, 4 μg/mL) antibodies, with germinal center counts determined by GL7+ cell clusters. Imaging was performed using an EVOS™ M7000 Imaging System (Thermo Fisher Scientific) with consistent exposure settings across experimental groups.

### SARS-CoV-2 pseudovirus production and neutralization assay

HEK293T cells were transfected at 60–70% confluency in 10-cm dishes using jetPRIME™ transfection reagent (Polyplus) with a plasmid mixture containing 10 μg psPAX2, 6 μg pLenti-Luci-GFP, and 4 μg pcDNA3.1-Spike (SARS-CoV-2 JN.1, WA1/D614G, B.1.617.2, or BA.2.86 strain) in DMEM (total 20 μg DNA). After 48 to 72 h, pseudovirus-containing supernatants were harvested, centrifuged (3,000 × g, 10 min, 4°C), aliquoted, and stored at −80°C until use.

SARS-CoV-2 neutralizing antibody titers were tested as described previously with slight modifications(*69*). Briefly, HEK293T-hACE2 cells were seeded in 96-well plates (4×10□ cells/well) and cultured for 24 h to reach >90% confluency. Serially diluted serum samples were mixed with SARS-CoV-2 pseudovirus and incubated at 37°C for 1 h before transfer to the pre-seeded plates. Following a 21–24 h incubation at 37°C with 5% CO□ (with DMEM as a negative control), supernatants were aspirated and replaced with luciferase substrate (Cat. No.11404ES60, YEASEN). Luminescence was quantified after a 2-minute dark incubation using a Spark® multimode microplate reader (Tecan). The 50% pseudovirus neutralization titer (pVNT_50_) was defined as the serum dilution achieving ≥50% reduction in relative luminescence units (RLU) compared to virus-only controls.

### Fluorescent antibody virus neutralization (FAVN) assay

Rabies virus neutralizing antibody (VNA) titers were quantified using a fluorescent antibody virus neutralization (FAVN) assay as previously described(*20*). Briefly, 100 μL of DMEM was added to a 96-well plate, and 50 μL of serum or standard serum was added to the first column in quadruplicate and subjected to threefold serial dilutions (1:3) across subsequent columns. A 50 μL suspension containing 100 focus-forming units (FFU) of rabies challenge virus strain CVS-11 was added to each well. After 1-hour incubation at 37°C, 2 × 10□ BSR cells were seeded per well and cultured for 72 h at 37°C (5% CO□). Cells were fixed with 80% (v/v) ice-cold acetone for 30 min at −20°C and stained with a fluorescein isothiocyanate (FITC)-conjugated anti-rabies virus nucleoprotein (RABV N) monoclonal antibody (Cat. No. 800-092, Fujirebio) for 1 h at 37°C. Fluorescent foci were visualized using an Olympus IX51 epifluorescence microscope (Olympus, Tokyo, Japan). Neutralization titers were calculated by comparing fluorescence intensities to the NIBSC reference serum standard. Results were normalized and expressed in International Units per milliliter (IU/mL), defined as the reciprocal serum dilution reducing FFU counts by ≥50% relative to virus-only controls.

### Transcriptomic profiling of Pan-B cell isolated from inguinal lymph node

Transcriptomic profiling was performed on pan-B cells isolated from iLNs of C57BL/6 mice (female, 7–8 weeks old) intramuscularly immunized with 10□ TCID□□ Ad5-HA_PR8_ or Ad5-HA_PR8_-VLP in 100 μL PBS. At 7 dpi, iLNs (n=5 per group) were aseptically excised and mechanically dissociated through a 70-μm cell strainer to generate single-cell suspensions. Untouched pan-B cells were isolated via negative selection using the MojoSort™ Mouse Pan-B Cell Isolation Kit (Cat. No. 480052, BioLegend), with cell purity (>90%) confirmed by flow cytometry (FCM) (anti-CD45, anti-CD3, and anti-B220 antibodies; BioLegend). Total RNA was extracted from 1×10□ sorted cells using TRIzol® Reagent (Invitrogen) followed by DNase I treatment (Cat. No. AM2222, Thermo Fisher) to eliminate genomic DNA. RNA integrity was verified using an Agilent Bioanalyzer 2100 (RNA Integrity Number [RIN] >8.0). Stranded mRNA sequencing libraries were prepared from 1 μg total RNA via the Illumina TruSeq Stranded mRNA Library Prep Kit (poly-A selection), followed by paired-end sequencing (150 bp) on an Illumina NovaSeq 6000 platform (40 million reads/sample). Raw reads were quality-filtered using Trimmomatic (v0.39; SLIDINGWINDOW:4:20, MINLEN:36) and aligned to the mm10 reference genome with STAR aligner (v2.7.10a; outSAMtype BAM SortedByCoordinate). Differentially expressed genes (DEGs) were identified using DESeq2 (v1.38.3; adjusted p <0.05, |log_2_ (fold change)| >1), followed by Gene Ontology (GO) and Kyoto Encyclopedia of Genes and Genomes (KEGG) pathway enrichment analyses via clusterProfiler (v4.0.5; hypergeometric test, Benjamini-Hochberg correction). Sequencing services were provided by Wuhan MetWare Biotechnology Co., Ltd.

### Quantitative real-time PCR (qRT-PCR)

In this study, gene expression analysis was performed using qRT-PCR following standardized protocols. Reverse transcription of Pan B cell transcriptome RNA subsets was conducted with HiScript II^®^ Reverse Transcriptase (Vazyme) using oligo(dT) primers, followed by qPCR amplification in 384-well plates on an ABI PRISM^®^ 7900HT system (Applied Biosystems). Reactions contained SYBR Green Master Mix (Vazyme), 10 μM gene-specific primers (Table.S2), and 1 μL cDNA template, with thermal cycling parameters: 95°C for 3 min, 40 cycles of 95°C for 10 sec and 60°C for 30 sec, followed by melt curve analysis. The 2×-ΔΔCt method was employed for relative quantification using *GAPDH* as the reference gene.

### Influenza viral plaque assay

Lungs harvested from infected mice were homogenized in DMEM (1 mL per organ) and stored at −80°C. For plaque quantification, MDCK cells were seeded in 12-well plates (1.5 × 10□ cells per well) 24 h prior to infection. Confluent monolayers were washed with PBS and inoculated with 1 mL of serially diluted lung homogenates. After 1 h of adsorption at 37°C, 5% CO□, with gentle agitation at 15-min intervals, the inoculum was replaced with an overlay medium containing DMEM, 1.6% low-melting-point agarose, and 1 μg/mL TPCK-trypsin (Sigma-Aldrich). Plates were incubated for 72 h at 37°C, 5% CO□, fixed with 4% (v/v) paraformaldehyde, and stained with 0.5% (w/v) crystal violet. Viral plaques were counted manually, and titers were calculated as plaque-forming units per milliliter (PFU/mL) using dilution-adjusted counts.

### Histopathological assessment

Following intranasal challenge with influenza virus PR8, lung tissues were collected from euthanized mice at predetermined time points, fixed in 4% paraformaldehyde for 48 h at 4°C, and processed through graded ethanol dehydration and paraffin embedding. Serial sections of 4–5 μm thickness were cut using a rotary microtome and mounted onto glass slides. Hematoxylin and eosin (H&E) staining was performed according to standard protocols: sections were deparaffinized, rehydrated, stained with Harris hematoxylin for 5 min, differentiated in acid-alcohol (1% HCl in 70% ethanol), counterstained with eosin Y for 1 min, and dehydrated through a graded alcohol series (70% to 100%) before cover-slipping with neutral balsam. Histopathological evaluation was conducted under double-blind conditions using bright-field microscopy to assess characteristic features, including interstitial pneumonia severity, inflammatory cell infiltration, alveolar wall thickening, hemorrhage, and epithelial damage. Representative images were captured at magnifications of 100×, 200×, and 400× for comparative analysis.

### *In vivo* imaging test

Following intranasal or intramuscular immunization with 10^7^ TCID_50_ Ad5-Luci, mice were anesthetized with isoflurane and intraperitoneally injected with 150 mg/kg D-luciferin potassium salt 2 min via intraperitoneal injection. Bioluminescence imaging was initiated 2 min post-injection using an IVIS^®^ Spectrum imaging system (PerkinElmer, MA, USA). Imaging was performed at specified timepoints post-immunization with optimized exposure times (1–60 s) to avoid pixel saturation. Total photon flux (photons s□¹ cm□² sr□¹) was quantified within anatomically defined regions of interest (ROIs: lungs, and injection site) using Living Image^®^ Software v4.7.3 (PerkinElmer).

### Statistical analysis

All statistical analyses were conducted using GraphPad Prism® v8.0 (GraphPad Software), with parametric tests selected based on experimental design. Survival analysis was performed using the Mantel-Cox log-rank test to compare statistical differences between Kaplan-Meier survival curves. One-way ANOVA (for single-variable comparisons) and two-way ANOVA (for multi-variable comparisons) were applied to evaluate intergroup differences in quantitative datasets, followed by Tukey’s post hoc test for pairwise comparisons where applicable. All graphical data display error bars representing standard deviation (SD) to quantify variability across biological replicates (n ≥ 3). Statistical significance thresholds were defined as follows: \**P* < 0.05, \*\**P* < 0.01, \*\*\**P* < 0.001, \*\*\*\**P* < 0.0001, with *P*-values adjusted for multiple comparisons using the Benjamini-Hochberg method where appropriate.

## Acknowledgments

We gratefully acknowledge the technical support from OE Biotech Co., Ltd (Shanghai, China), with special thanks to Dr. Wang Qing for her expert guidance and critical contributions to scRNA-seq data analysis. We are particularly grateful to Dr. Chen Jing from the College of Foreign Languages, Huazhong Agricultural University for her editing and revision of the manuscript. We would like to thank Xiao Shuang and Xu Yingying from the Public Instrument Center of the College of Animal Science and Technology and the College of Veterinary Medicine of Huazhong Agricultural University for their help in our FCM and transmission electron microscopy experiments.

## Data and materials availability

All data are present within the manuscript and Supplementary Data.

