## Supplementary Text Figures. S1 to S8 Tables S1 to 2 for "A chimeric Ad5-Envp-VLP vaccine platform confers broad-spectrum immunity against emerging and re-emerging pathogens"

**This PDF file includes:**

Supplementary Text

Figures. S1 to S8

Tables S1 to 2

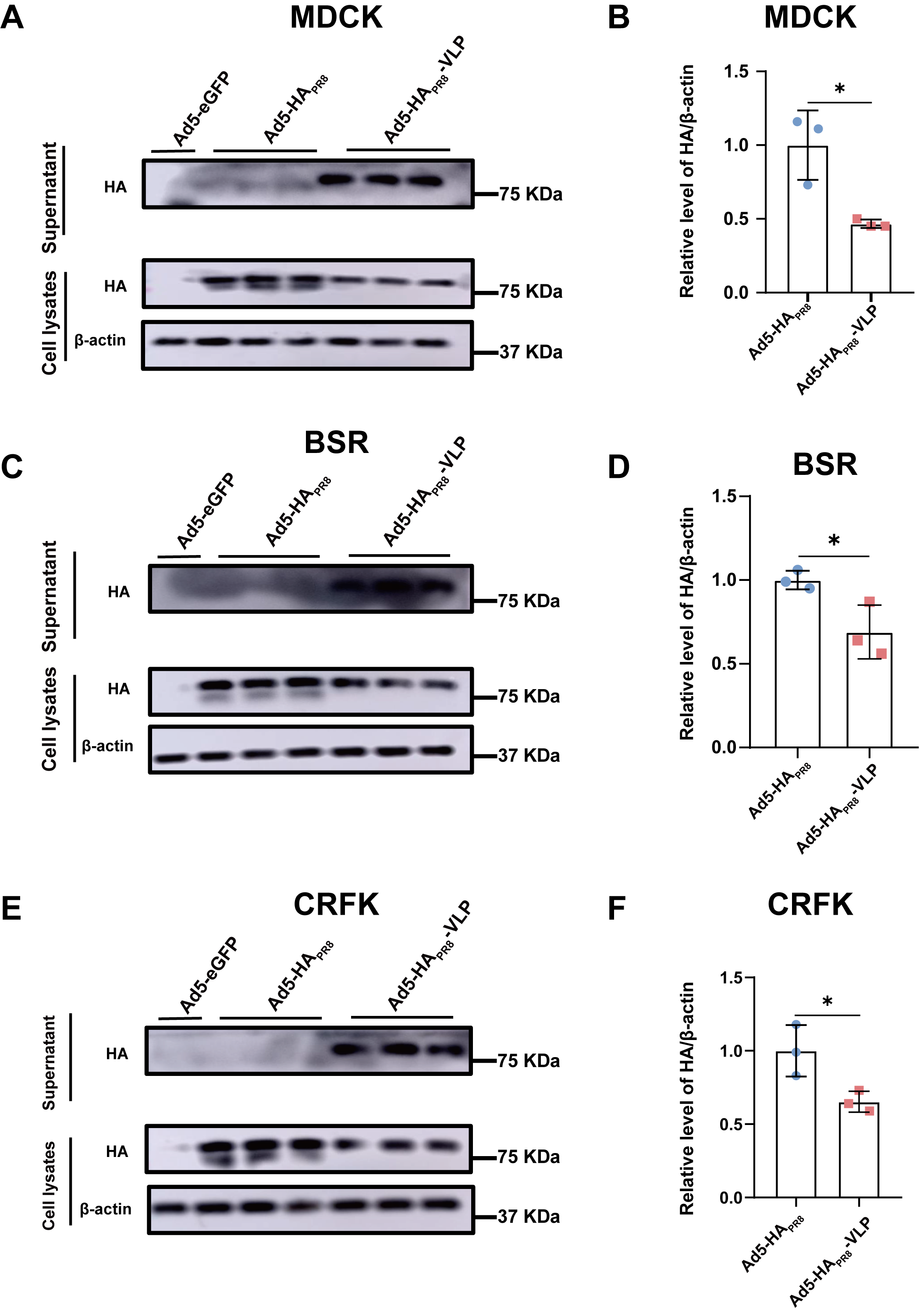

**Figure S1. Ad5-HA_PR8_-VLP mediates the budding of HA protein in multiple species.** (A-B) MDCK (canine), (C-D) BSR (hamster), and (E-F) CRFK (feline) cells infected with Ad5-HA_PR8_-VLP, Ad5-HA_PR8_, or Ad5-eGFP (MOI=1). Left panels: Representative Western blots of intracellular HA (lysates) and secreted HA (supernatants) at 48 hpi. Right panels: Quantification of intracellular HA (normalized to β-actin). Data are presented as the mean ± SD, and an unpaired two-tailed t-test was used in B, D and F. Western blotting data are the representative results of two independent experiments. NS: not significant; **P* < 0.05, ***P* < 0.01, ****P* < 0.005, *****P* < 0.0001.

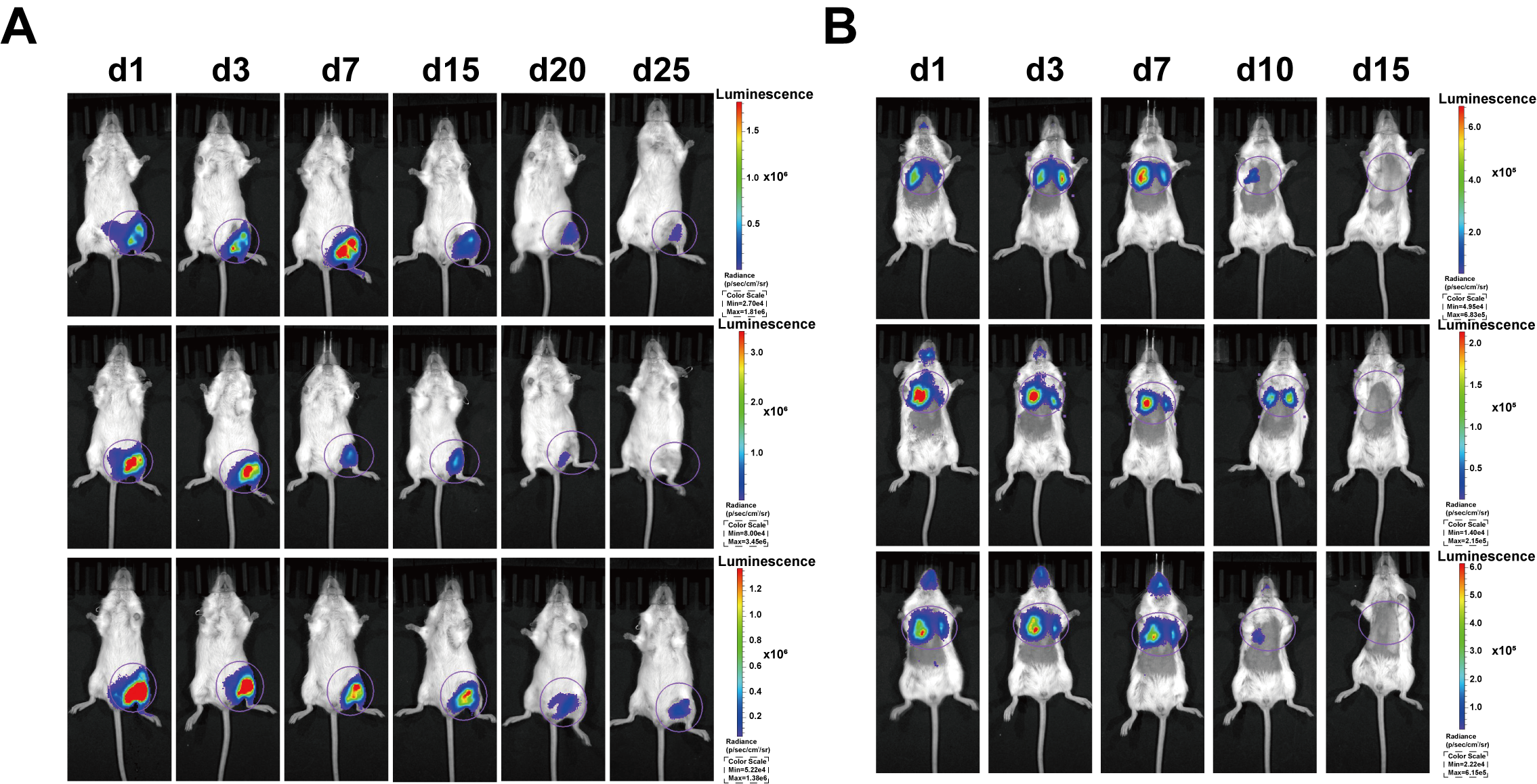

**Figure S2. Exogenous gene expression kinetics of Ad5-Luci following intramuscular or intranasal administration.** (A) BALB/c mice (n=3 per group) were injected intramuscularly with 10^7^ TCID_50_ Ad5-Luci. Longitudinal luciferase (Luci) expression in the injection site was quantified by IVIS imaging at day 1, 3, 7, 15, 20 and 25. (B) Mice (n=3 per group) received intranasal instillation of 10^7^ TCID_50_ Ad5-Luci, with Luci expression monitored in the lungs at day 1, 3, 7, 10 and 15.

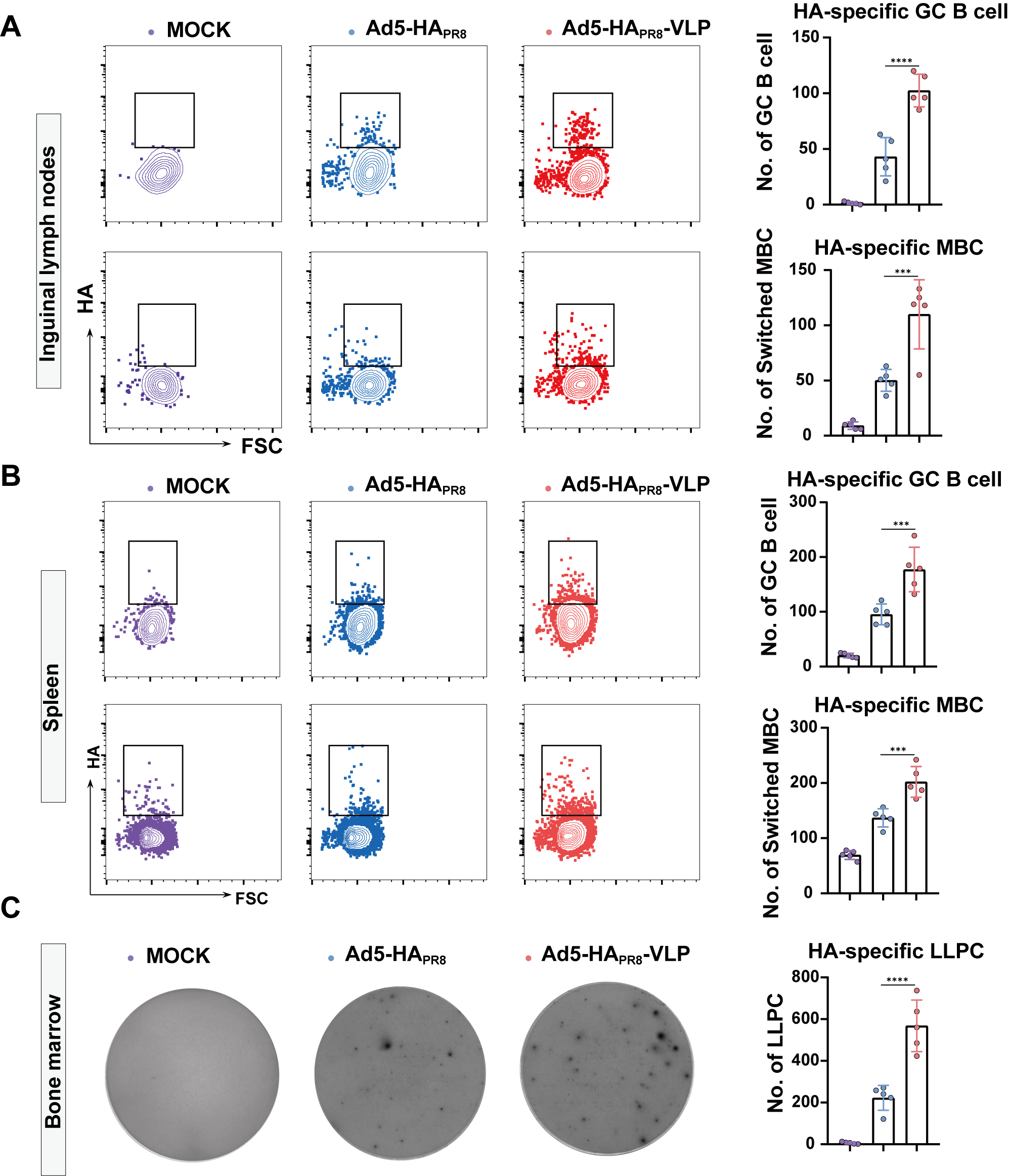

**Figure S3. Ad5-HA_PR8_-VLP immunization enhances HA-specific B cell responses.** C57BL/6 mice (n=5 per group) were intramuscularly immunized with Ad5-HA_PR8_-VLP or Ad5-HA_PR8_ (control). At 35 days post-immunization, inguinal lymph nodes (iLNs), spleen, and bone marrow were harvested for quantification of HA-specific B cell subsets. (A) Left panels show representative flow cytometry plots of HA-specific GC B cells (CD45+B220+CD3-CD95+GL7+HA+) and class-switched memory B cells (MBCs; CD45+B220+CD3-IgD-CD38+HA+) in iLNs; right panels display statistical quantification of these populations. (B) Left panels depict HA-specific GC B cells and MBCs in spleen via flow cytometry; Right panels showed corresponding quantitative results. (C) Left panels present representative ELISpot images of HA-specific LLPCs in bone marrow; right panels provided quantification of LLPCs. Data are presented as mean ± SD. Statistical significance was determined by one-way ANOVA with Tukey’s multiple-comparison test, with significance levels denoted as NS (not significant), **P* < 0.05, ***P* < 0.01, ****P* < 0.005, and *****P* < 0.0001.

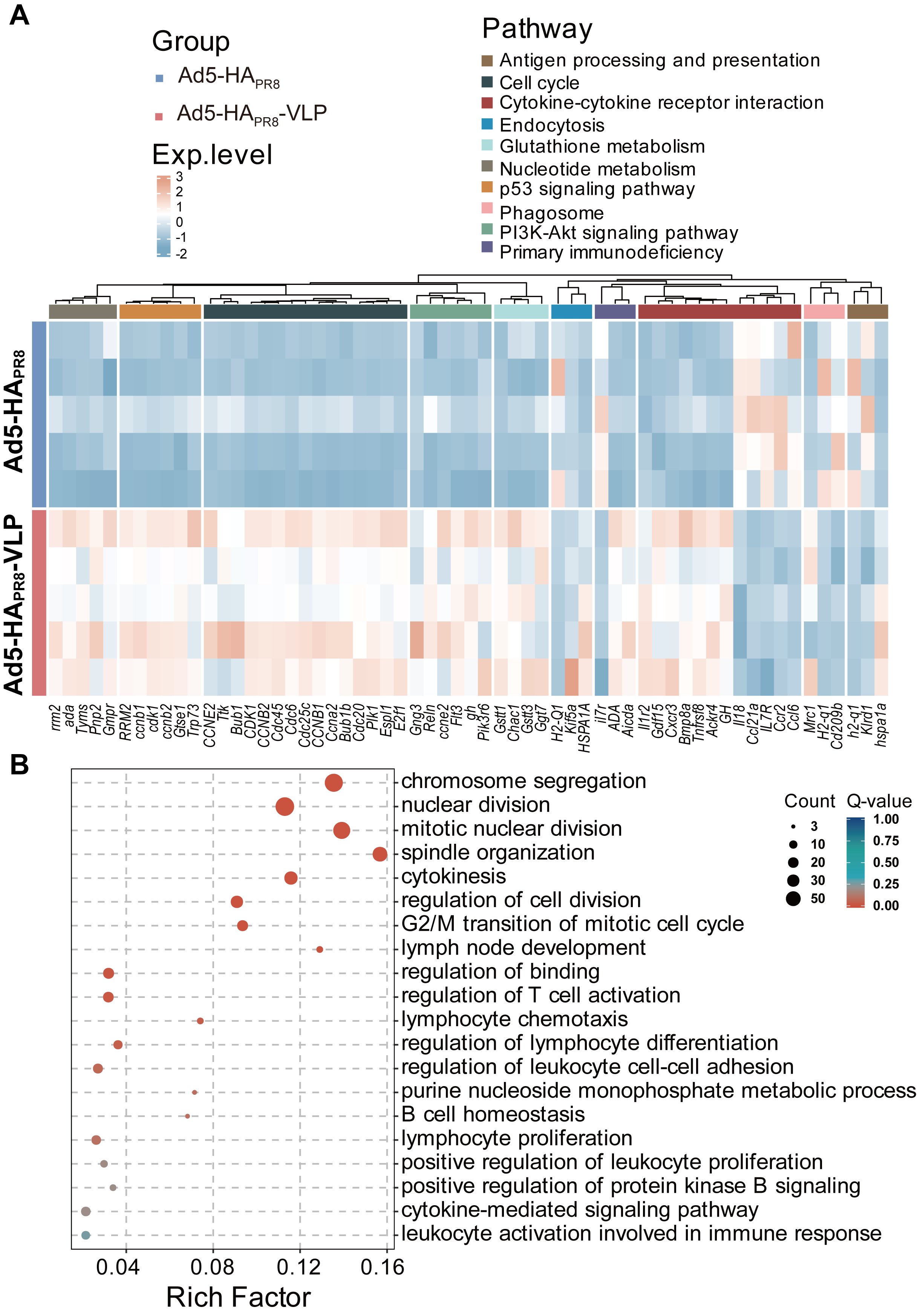
 **Figure S4. KEGG and GO enrichment analysis of differentially expressed genes in pan-B cells.** (A) Heatmap visualization of scaled gene expression levels for selected pathways of interest. (B) Top 20 GO term enrichment of DEGs.

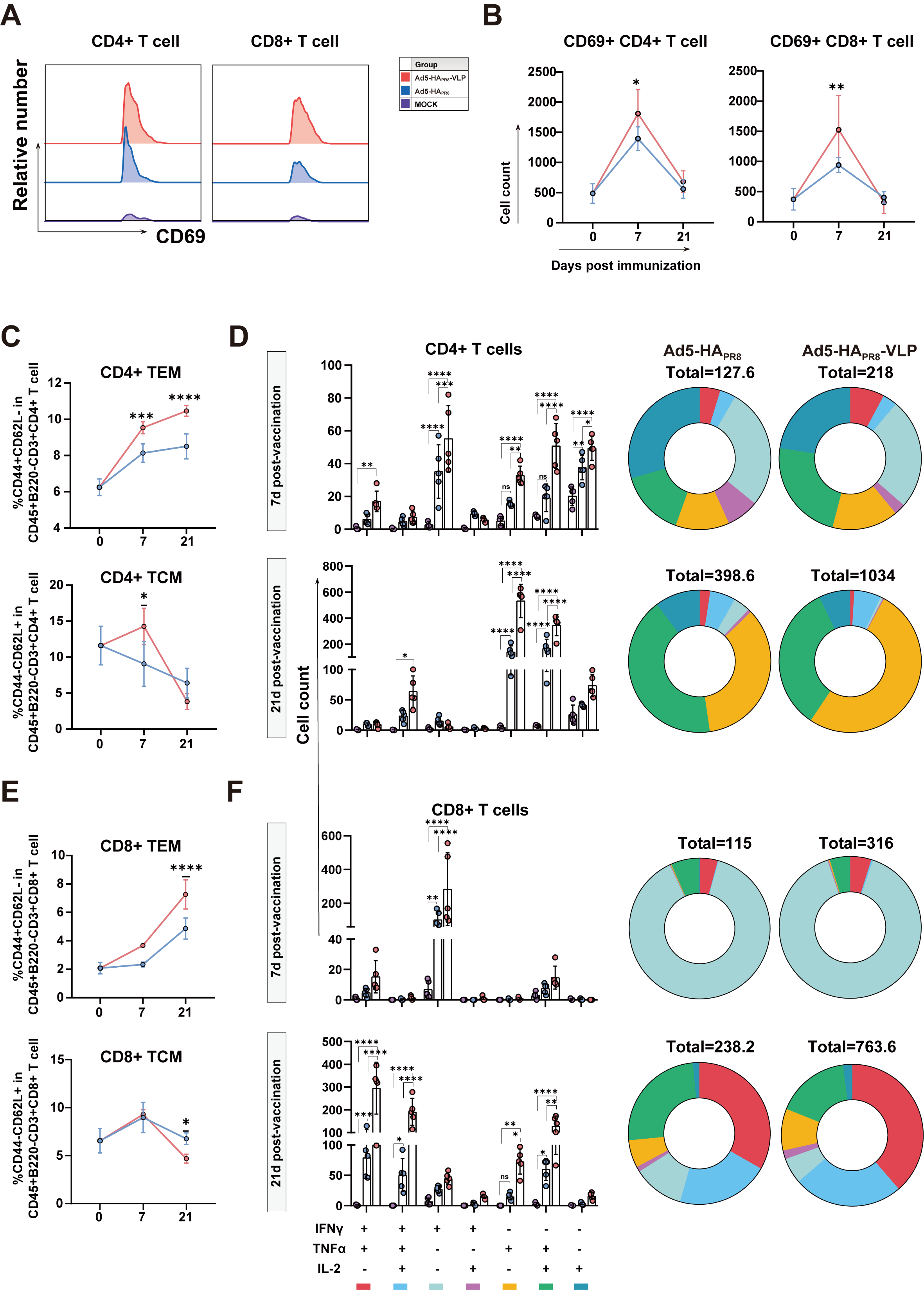

**Figure S5. T cell immune responses in Ad5-HA_PR8_-VLP-immunized mice.** C57BL/6 mice (n=5 per group) were intramuscularly immunized with Ad5-HA_PR8_-VLP or Ad5-HA_PR8_. Spleens were harvested at days 7 and 21 post-immunization to quantify HA-specific T cell responses. (A) Representative flow cytometry plots of CD69 expression in splenic CD4+ and CD8+ T cells at 7 dpi. (B) Quantification of CD69+ cells among CD4+ (left) and CD8+ T cells (right). (C) Proportions of CD4+ effector memory T cells (TEM; CD44+CD62L-) and central memory T cells (TCM; CD44+CD62L+) in spleens at day 7 (top) and 21 (bottom) (gating: CD45+B220-CD3+CD8-CD4+). (D) Quantification of cytokine-producing CD4+ T cells (IFN-γ+, IL-2+, TNF-α+) in spleens at day7 (top) and 21 (bottom). (E) Proportions of CD8+ TEM (CD44+CD62L-) and TCM (CD44+CD62L+) in spleens at day 7 (top) and 21 (bottom) (gating: CD45+B220-CD3+CD8+CD4-). (F) Frequencies of cytokine-producing CD8+ T cells (IFN-γ+, IL-2+, TNF-α+) in spleens at day 7 (top) and 21 (bottom). Pie charts illustrate the distribution of cells producing one, two, or three cytokines. Data are presented as mean ± SD. Statistical significance was determined by two-way ANOVA with Tukey’s multiple-comparison test; significance levels: NS (not significant), **P* < 0.05, ***P* < 0.01, ****P* < 0.005, and*****P* < 0.0001.

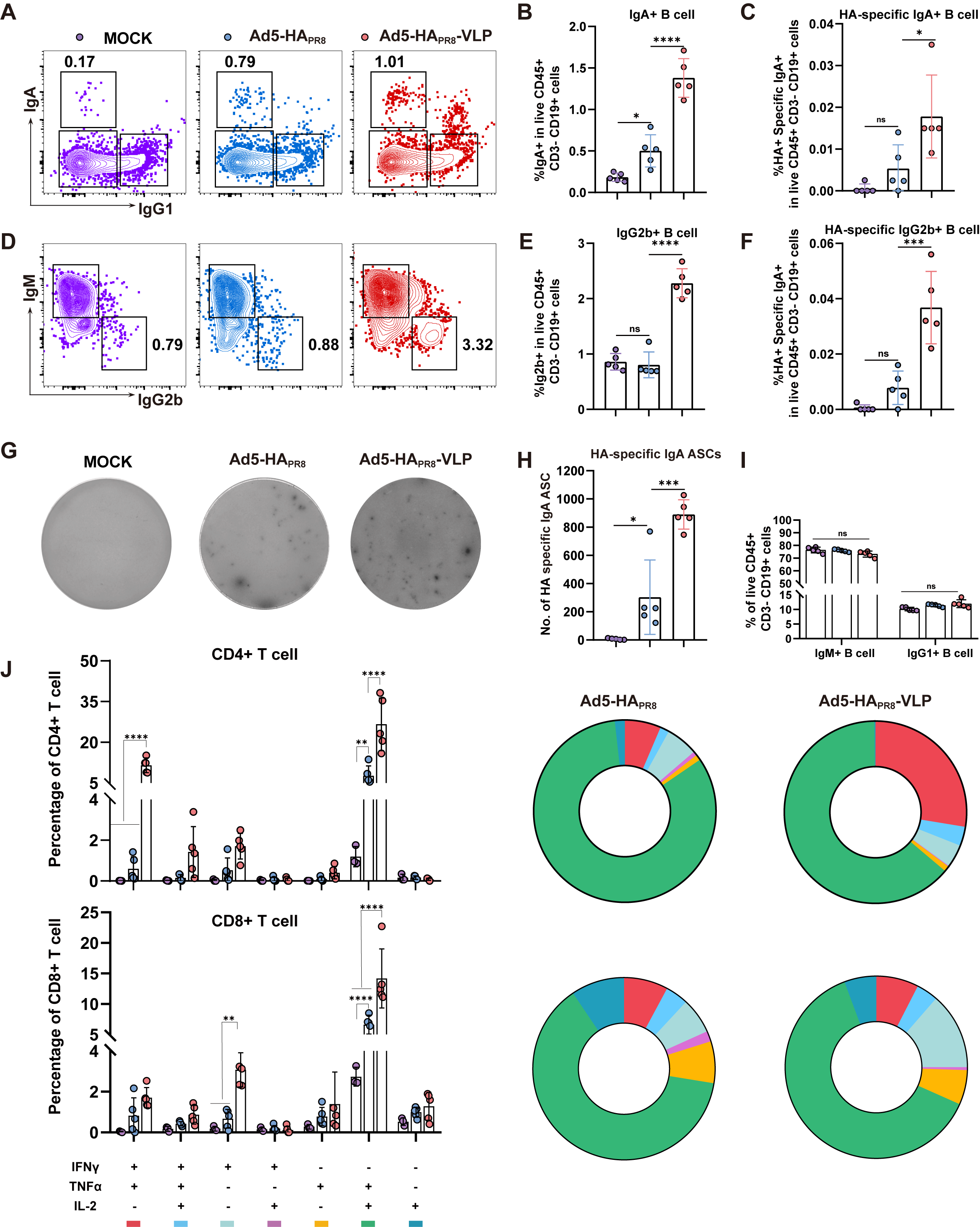

**Figure S6. VLPs formulation enhances HA-specific B and T cell responses in lungs.** C57BL/6 mice (n=5 per group) were intranasally immunized with 10⁷ TCID_50_ Ad5-HA_PR8_-VLP or Ad5-HA_PR8_. Lungs were harvested at day 21 for analysis of HA-specific B and T cell subsets. (A) Representative flow cytometry plots of IgA+ and IgG1+ B cells (CD45+CD19+CD3-) in lungs. (B, C) Proportions of total and HA-specific IgA+ B cells. (D) Representative flow cytometry plots of IgM+ and IgG2b+ B cells. (E, F) Proportions of total and HA-specific IgG2b+ B cells. (G) Representative ELISpot images of HA-specific IgA ASCs. (H, I) Quantification of HA-specific IgA ASCs and proportions of total IgM+ and IgG1+ B cells. (J) Frequencies of cytokine-producing CD4+ (IFN-γ+, IL-2+, TNF-α+) and CD8+ T cells in lungs. Pie charts show polyfunctional cytokine profiles (cells producing 1, 2, or 3 cytokines). Data are presented as mean ± SD. Statistical significance (Panels B, C, E, F, H and I) was determined by one-way ANOVA with Tukey’s multiple-comparison test; Panel J was analyzed by two-way ANOVA with Tukey’s test. Significance levels: NS (not significant), **P* < 0.05, ***P* < 0.01, ****P* < 0.005, *****P* < 0.0001.

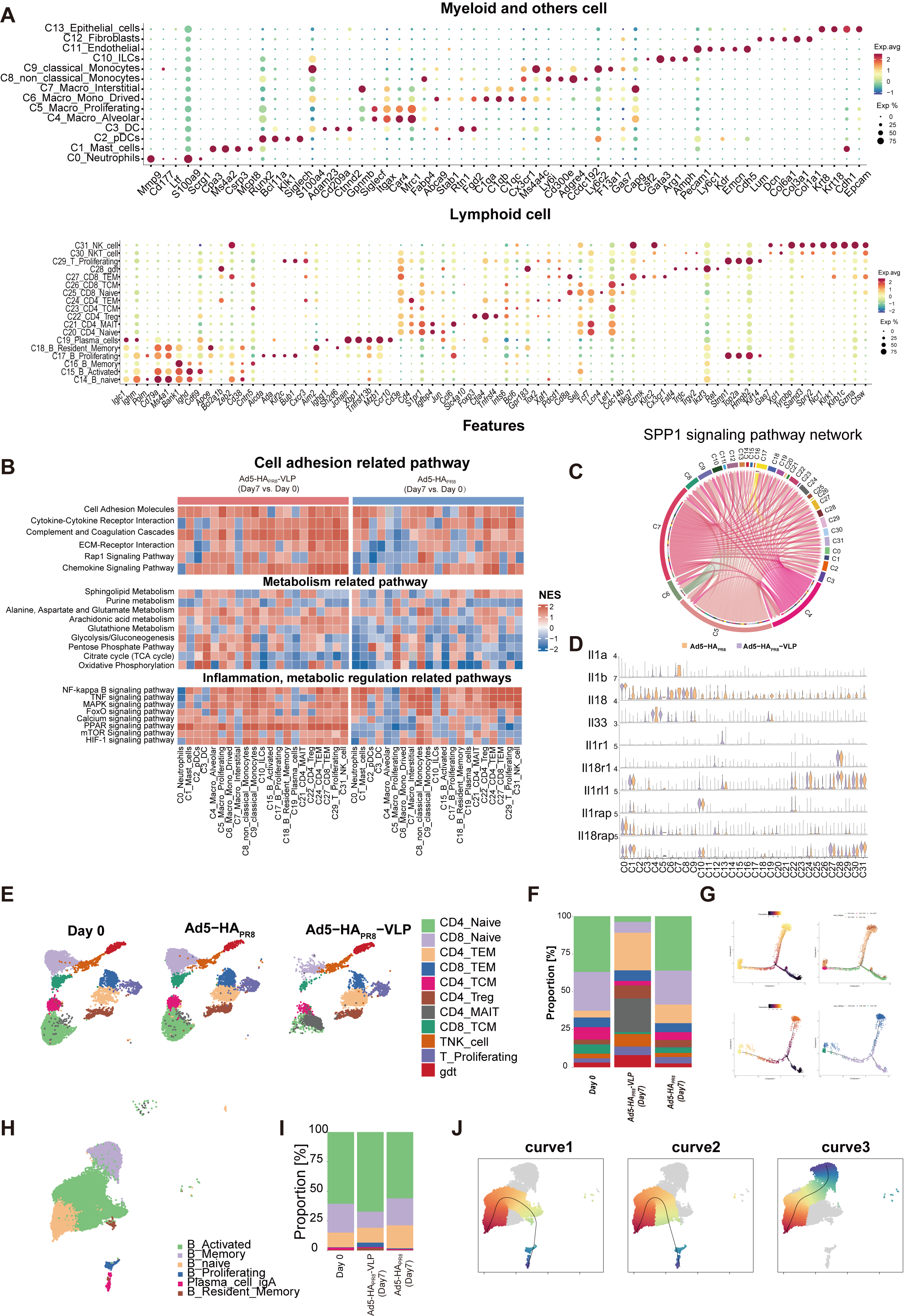

**Figure S7. scRNA-seq analysis of immune response induced by Ad5-HA_PR8_-VLP and Ad5-HA_PR8_.** (A) Clusters and their associated cluster-specific genes. (B) Significantly enriched *Cell adhesion*, *Metabolism*, *Inflammation*, *metabolic regulation* related KEGG pathways across clusters (C0-1, C15, C17, C18, C19, C21, C22, C24, C27, C29 and C31) at day 7. Only clusters with a significantly modulated pathway are shown. (C) Chord diagram showing SPP1 signaling network. The edge color and arrow represent sender source and edge weights indicate interaction strength represented by computed communication probability. (D) Violin plot of expression distribution of genes related to IL1 signaling pathway. (E) UMAP of subclusters in T cell. (F) Proportion of T cell subsets in different immune groups. (G) Left: the pseudo-time trajectory of T cells, the colors change from dark to light, indicating the time of differentiation from early to late stages. Right: the pseudo-time trajectory of T cell population differentiation, each color represents a cell population. (H) UMAP of B cell types clustered by single-cell transcriptional analysis. (I) Proportion of B cell subsets in different immune groups. (J) The schematic illustrates the hierarchical structure and sequential progression of B-cell lineage differentiation. Each point represents an individual cell, with the solid black line delineating the differentiation trajectory. A color gradient from red to blue indicates advancing pseudotime (early to late stages).

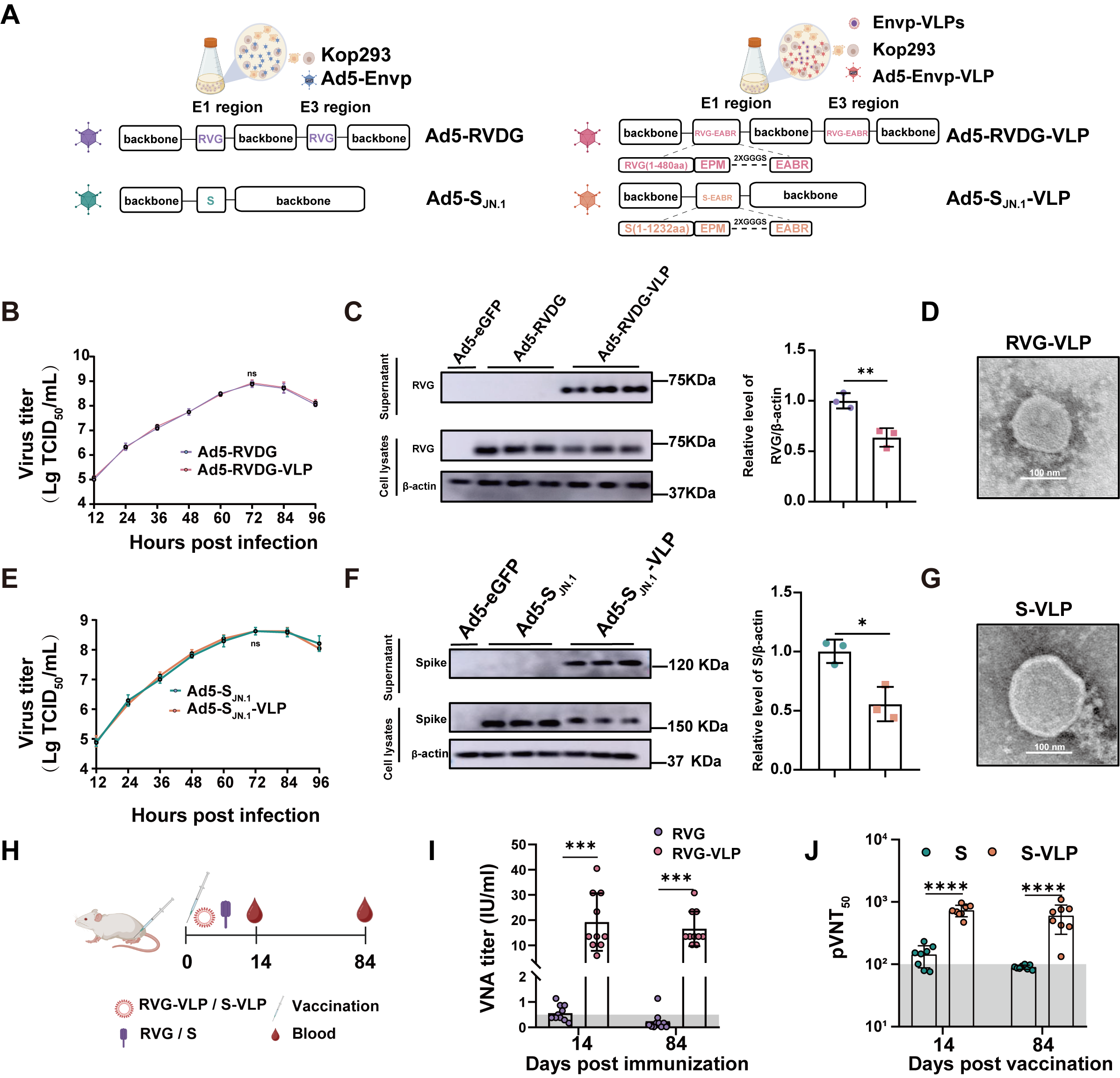

**Figure S8. Construction and characterization of Ad5-S_JN.1_-VLP and Ad5-RVDG-VLP.** (A) Schematic design of Ad5-Envp and Ad5-Envp-VLP. The EABR motif was fused to cytoplasmic domain-truncated RVG (1–480) and SARS-CoV-2 S (1–1232). Parental adenoviruses Ad5-RVDG and Ad5-S_JN.1_ encoding full-length proteins served as controls. (B, E) Replication kinetics of Ad5-Envp-VLP (EABR-modified) and Ad5-Envp (full-length) in HEK 293 cells (n=3). (C, F) Western blot analysis of envelope protein expression in cell lysates and supernatants from Ad5-RVDG-VLP- and Ad5-S_JN.1_-VLP-infected cells versus controls. Transmission electron microscopy of SEC-purified VLPs from Ad5-RVDG-VLP- (D), and Ad5-S_JN.1_-VLP- (G) infected cells, revealing spherical particles (90–110 nm) with surface projections. Scale bars: 100 nm. (H) Mice were immunized intramuscularly with DMEM, RVG-VLP, RVG, S-VLP, or soluble S (AS03-adjuvanted), respectively. Virus-neutralizing antibody (VNA) titers against RABV (I) and SARS-CoV-2 (J) in sera. Data are mean ± SD. Statistical analysis: unpaired two-tailed t-test (C, F) and two-way ANOVA with Tukey’s test (I, J). NS: not significant; **P* < 0.05, ***P* < 0.01, ****P* < 0.005, *****P* < 0.0001.

**Table. S1** **Pairs of backbone and shuttle plasmids**

| **backbone plasmid** | **shuttle plasmid** | **recombinant** **adenoviruses** |
| --- | --- | --- |
| pBHGcre/loxp | pDC315-S | Ad5-S_JN.1_ |
| pBHGcre/loxp | pDC315-S-EABR | Ad5-S_JN.1_-VLP |
| pBHGcre/loxp-RVG | pDC315-RVG | Ad5-RVDG |
| pBHGcre/loxp-RVG-EABR | pDC315-RVG-EABR | Ad5-RVDG-VLP |
| pBHGcre/loxp-HA | pDC315-HA | Ad5-HA_PR8_ |
| pBHGcre/loxp-HA-EABR | pDC315-HA-EABR | Ad5-HA_PR8_-VLP |

**Table.S2** **Primers used for qPCR**

| **Primer name** | **Sequence (5’ to 3’)** |
| --- | --- |
| Mus-GAPDH-F | AGGTCGGTGTGAACGGATTTG |
| Mus-GAPDH-R | TGTAGACCATGTAGTTGAGGTCA |
| Mus-Aicda-F | GCCACCTTCGCAACAAGTCT |
| Mus-Aicda-R | CCGGGCACAGTCATAGCAC |
| Mus-Ada-F | ACCCGCATTCAACAAACCCA |
| Mus-Ada-R | AGGGCGATGCCTCTCTTCT |
| Mus-Aurka-F | CTGGATGCTGCAAACGGATAG |
| Mus-Aurka-R | CGAAGGGAACAGTGGTCTTAACA |
| Mus-Aurkb-F | CAGAAGGAGAACGCCTACCC |
| Mus-Aurkb-R | GAGAGCAAGCGCAGATGTC |
| Mus-Cdk1-F | AGAAGGTACTTACGGTGTGGT |
| Mus-Cdk1-R | GAGAGATTTCCCGAATTGCAGT |
| Mus-Cdc45-F | GAGGTTCCTGCCTACGACG |
| Mus-Cdc45-R | TCCTGTTTCGCTCCACTATCT |
| Mus-Ccne2-F | ATGTCAAGACGCAGCCGTTTA |
| Mus-Ccne2-R | GCTGATTCCTCCAGACAGTACA |
| Mus-Cdc20-F | TTCGTGTTCGAGAGCGATTTG |
| Mus-Cdc20-R | ACCTTGGAACTAGATTTGCCAG |
| Mus-Ccnb1-F | AGAGCTATCCTCATTGACTGGC |
| Mus-Ccnb1-R | AACATGGCCGTTACACCGAC |
| Mus-Ccnb2-F | GCCAAGAGCCATGTGACTATC |
| Mus-Ccnb2-R | CAGAGCTGGTACTTTGGTGTTC |
| Mus-Pik3r6-F | AGCAATCAGGGCATGTGGAG |
| Mus-Pik3r6-R | CGTCCGTCCTCGCTTTCTG |
| Mus-Stmn1-F | TCTGTCCCCGATTTCCCCC |
| Mus-Stmn1-R | AGCTGCTTCAAGACTTCCGC |
| Mus-IL7R-F | GCGGACGATCACTCCTTCTG |
| Mus-IL7R-R | AGCCCCACATATTTGAAATTCCA |
